# Neural Regression, Representational Similarity, Model Zoology & Neural Taskonomy at Scale in Rodent Visual Cortex

**DOI:** 10.1101/2021.06.18.448431

**Authors:** Colin Conwell, David Mayo, Michael A. Buice, Boris Katz, George A. Alvarez, Andrei Barbu

## Abstract

How well do deep neural networks fare as models of mouse visual cortex? A majority of research to date suggests results far more mixed than those produced in the modeling of primate visual cortex. Here, we perform a large-scale benchmarking of dozens of deep neural network models in mouse visual cortex with both representational similarity analysis and neural regression. Using the Allen Brain Observatory’s 2-photon calcium-imaging dataset of activity in over 6,000 reliable rodent visual cortical neurons recorded in response to natural scenes, we replicate previous findings and resolve previous discrepancies, ultimately demonstrating that modern neural networks can in fact be used to explain activity in the mouse visual cortex to a more reasonable degree than previously suggested. Using our benchmark as an atlas, we offer preliminary answers to overarching questions about levels of analysis (e.g. do models that better predict the representations of individual neurons also predict representational similarity across neural populations?); questions about the properties of models that best predict the visual system overall (e.g. is convolution or category-supervision necessary to better predict neural activity?); and questions about the mapping between biological and artificial representations (e.g. does the information processing hierarchy in deep nets match the anatomical hierarchy of mouse visual cortex?). Along the way, we catalogue a number of models (including vision transformers, MLP-Mixers, normalization free networks, Taskonomy encoders and self-supervised models) outside the traditional circuit of convolutional object recognition. Taken together, our results provide a reference point for future ventures in the deep neural network modeling of mouse visual cortex, hinting at novel combinations of mapping method, architecture, and task to more fully characterize the computational motifs of visual representation in a species so central to neuroscience, but with a perceptual physiology and ecology markedly different from the ones we study in primates.

## 1 Introduction

To date, the most successful models of biological visual cortex are object-recognizing deep neural networks applied to the prediction of neural activity in primate visual cortex [1–5]. Corresponding to the biology not only at the level of individual layers, but across the feature hierarchy, these models are so powerful they can now effectively be used as neural controllers, synthesizing stimuli that drive neural activity far beyond the range evoked by any handmade experimental stimulus [6]. The correspondence of these same models to mouse visual cortex, on the other hand, has proven a bit more tenuous [7, 8], with a recent finding even suggesting that randomly initialized networks are as predictive of rodent visual cortical activity as trained ones [9].

Often implicit in interpretations of these results is the notion that the visual milieu and machinery of mice is simply *different* – something characterized more, perhaps, by brute force predator avoidance and the ‘flexible random associations’ thought to define senses like olfaction [10] than by the sophisticated active sampling and representational compositionality enabled by primate central vision. And yet, mice do recognize objects [11, 12] – and do engage in other sophisticated visual behaviors [13] that suggest they must have visual solutions that at least functionally approximate the kinds of solutions learned by modern computer vision algorithms. If these models perform well in monkeys, but not in mice, are we overfitting to an artifact? Are the object recognition capabilities of mice simply the byproduct of a representational competence learned through other (even more behaviorally relevant) tasks? Have mice perhaps converged on solutions to visual problems that fundamentally differ from the solutions that undergird the emergent similarity between monkeys and machines? To even begin to answer these questions, we need substantially more comprehensive modeling statistics than we currently have. Our main goal in this work was to provide exactly that – to re-examine at large scale the state of neural network modeling in the visual cortices of mice, using many thousands of neurons, over 110 distinct neural network models, and two methods of mapping models to brain.

We summarize the statistics from our benchmarking survey in five main results:

1. Training matters. The randomly initialized variants of some convolutional architectures fare well when predicting individual neural responses, but representational similarity is always better captured by features learned in service of some task. (Segmentation seems best.)
2. Features of intermediate complexity dominate in the prediction of all cortical sites, but both our mapping methods do demonstrate an upwards gradient in complexity from primary visual cortex onwards that roughly matches the information processing hierarchy proposed elsewhere in the rodent neurophysiology literature.
3. Taskonomic tools that have previously been shown to approximate functional organization in primates fail to strongly differentiate anatomical regions in mice, with the same kinds of tasks dominant across multiple, distinct neural sites.
4. When aggregated in similar ways, representational similarity and neural regression methods capture similar trends in the kinds of feature spaces that best predict the biology.
5. While still far from the overall noise ceiling for this highly reliable neural data, a variety of the artificial deep net models in our survey make predictions only slightly less accurate than ‘biological conspecific models’ composed of the neurons from other mice.

## 2 Methods

### 2.1 Neural Dataset

For neural data, we use the Allen Brain Observatory Visual Coding^1^ dataset [14] collected with two-photon calcium-imaging from the visual cortex of 256 awake adult transgenic mice and consisting of approximately 59,610 unique, individual neurons. Calcium-imaging fluorescence patterns are preprocessed and deconvolved by the Allen Institute^2^. The neurons sampled include neurons from 6 visual cortical areas at 4 cortical depths across 12 genetic cre lines. The visual experiments recorded activity for both artificial images (e.g., diffraction gratings) and 118 natural scenes. We analyze only the latter to ensure comparable inputs to what is typically used in the training of deep nets. Each natural scene is displayed 50 times over the course of an assay.

To ensure an optimal signal to noise ratio, we perform a significant amount of subsetting on the full neural population, beginning by subsetting only excitatory neurons. Recent analyses suggest neural activity throughout mouse visual cortex is often impacted by extraneous, external body movements [15]. For this reason, we subsequently filter out any neurons whose peak responses to the presentation of natural scene images are significantly modulated by the mouse’s running speed, using an ANOVA metric provided by the Allen Institute. We further subselect neurons by assessing their split-half reliability across trials (with each split-half constituting 25 of 50 presentations for each image), keeping only those neurons exhibiting 0.8 reliability and above. This thresholding still leaves 6619 neurons for analysis, is in line with prior work on primates, and supports, for example, the construction of cortical representational dissimilarity matrices (RDMs) with split-half reliabilities as high as 0.93. (More details on the relationship between our metrics and neural reliability, including visualizations of some of our results across many degrees of thresholding, can be found in A.4 of the Appendix.)

### 2.2 Model Zoology

To explore the influence of model architecture on predictive performance, we use 26 model architectures from the Torchvision (PyTorch) model zoo [16] and 65 model architectures from the Timm [17] model zoo [18–52]. These models include convolutional networks, vision transformers, normalization-free networks and MLP-Mixer models. For each of these models, we extract the features from one trained and one randomly initialized variant (using whatever initialization scheme the model authors deemed best) so as to better disentangle what training on object recognition affords us in terms of predictive power.

### 2.3 Neural Taskonomy

Model zoology provides decent perspective on the computations related to object recognition, but the responsibilities of the visual cortex (no matter the species) extend far beyond identifying the category of an object. To probe a wider range of tasks, we turn to Taskonomy: a single architecture trained on 24 different common computer vision tasks [53], ranging from autoencoding to edge detection. The model weights we use are from updated PyTorch implementations of the original Tensorflow models [54]. Key to the engineering of Taskonomy is the use of an encoder-decoder design in which only the construction of the decoder varies across tasks. While recent analyses using a similar approach in human visual cortex with fMRI data [55] have tended to focus only on the latent space of each task’s encoder, we extract representations *across all layers*, better situating Taskonomy within the same empirical paradigm that has so far defined the modeling of object recognition in the primate brain. For further clarity, we cluster the 24 tasks according to their ‘Taskonomic’ category — a total of 5 clusters (2D, 3D, semantic, geometric or other) that we further collapse into 4 clusters (lumping the only member of the ‘other’ category — a denoising autoencoder — in with its closest cousin — a vanilla autoencoder in the ‘2D’ category). These purely data-driven clusters are derived from estimates of how effectively a set of features learned for one task transfer to (or boost the performance in) another task [53]. Use of the Taskonomy models provides a unique opportunity to test variance in training regimes without the confound of simultaneous changes in architecture.

### 2.4 Self-Supervised Models

Full category supervision, while robust in its ability to build representations that transfer well to a variety of tasks, suffers in its neuroscientific relevance as an ethologically plausible mode of learning. Recently, self-supervised models have begun to provide viable alternatives to the representations learned by category-supervised models in both computer vision [56, 57] and neural mapping [58, 59]. Here, we assess 22 self-supervision models from the VISSL model zoo [60], ranging from earlier iterations (e.g. DeepCluster [61]) to modern contrastive learning algorithms (e.g. BarlowTwins and Dino [62–65]). We use these models to assess whether category-supervision, however powerful it is in predicting neural activity, might eventually be supplanted by these more realistic alternatives. 14 of these models have as their base architecture a standard ResNet50; 8 are built atop vision transformers.

### 2.5 Comparing Representations across Biological & Artificial Networks

Two methods predominate in the comparison of neural recordings to deep neural networks: at the most abstract level, one of these compares representational geometries computed across the activations of many individual neurons [66, 67]; the other attempts to predict the activity of individual neurons directly [67, 68]. Both of these techniques are grounded in the use of image-computable models and a shared stimulus set, but differ in the types of transformation applied to the neural activity generated by those stimuli. Given the difference in both target (neural populations versus individual neurons) and transforms (correlation matrices versus dimensionality reduction) we attempt a variant of each type of analysis here, comparing the two directly on the exact same neural data, with the same models and the same stimulus set, and in a granular, layer-by-layer fashion. (A more comprehensive review of neural mapping methods is provided in Section A.2 of the Appendix.)

#### 2.5.1 Representational Similarity Analysis

To compare the representational geometries of a given model to the representational geometries of the brain, we begin by computing classic representational dissimilarity matrices (RDMs) [69]. We compute these RDMS by calculating the pairwise correlation coefficients between the neural response vectors for each image (one for each of the 6 cortical areas surveyed). We then repeat this procedure for the artificial networks, aggregating the responses of the artificial neurons in a given layer, before aggregating them once more into a correlation matrix. We then measure the relationship between the RDMs computed from the biological and artificial networks with a second-order Pearson correlation between the flattened upper triangles of each. The resultant coefficient constitutes the score for how well a given model layer predicts the representational similarity of a given cortical area.

#### 2.5.2 Neural Regression (Encoding Models)

To more directly compare the biological and artificial neural activations in our data, we use a style of regression made popular in the modeling of primate visual cortex, epitomized by BrainScore [4]. Variants of this approach abound, but most consist of extracting model activations, performing dimensionality reduction, and then some form of cross-validated penalized or principal components regression. The dimensionality-reduced feature spaces of the model are used as the regressors of the activation patterns in a given neuron. After testing a number of these variants, we settled on sparse random projection for dimensionality reduction (which proved far more computationally efficient than standard PCA, without sacrifice in terms of regression scores), followed by ridge regression (in place of the more frequently used partial least squares regression).

The details of our method (programmed with [70]) are as follows: Given a network, we first extract a predetermined number of sparse random projections (4096, in this case) from the features of each layer — in line with the Johnson-Lindenstrauss lemma for the number of observations (images shown to the mice) in our data set ^3^. After extracting these projections, we regress them on the activity of each individual neuron using ridge regression (with a default lambda penalty of 1.0). The use of a penalized regression in this case allows us to monopolize generalized cross-validation (a linear algebraic form of leave-one-out cross-validation), yielding a set of predictions for the activity of each neuron for each image^4^. We then compute the Pearson correlation between the predicted and actual activity for each neuron to obtain a score per neuron per model layer, which we then aggregate by taking the mean of scores per neuron across cortical area.

We verify the efficacy of this method on the publicly available benchmarks of primate BrainScore, where (relative to BrainScore’s in-house regression method) we demonstrate provisional gains not only in terms of predictive score (sometimes up to *r* = 34%), but also in terms of speed and computational efficiency. (Details may be found in Section A.1 of the Appendix.)

### 2.6 Model Rankings

To rank the models according to how well they predict the variance in a given cortical area, we take the max across layers. In effect, this requires that a model ‘commit’ only one layer to the prediction of each area. In the case of our neural regression metric we call these scores the ‘SRP-Ridge Max’; in the case of our representational similarity metric we call these scores the ‘RSA Max’. A final mean taken over the SRP-Ridge Max and RSA Max scores per model per cortical area yields our overall model rankings, which serve as the basis for the bulk of our analyses.

### 2.7 Non-Neural Network Baselines

Prior to the ascendancy of neural network models, a significant amount of time and craft was invested in the hand-engineering of features to simultaneously facilitate image recognition and capture meaningful subsets of neural variance. In this work, we test how well a small subset of those features are able to explain the variance in rodent visual cortex, using both our neural encoding and representational similarity metrics. Our non-neural network baselines consist of random fourier features [71] (computed specifically to match the dimensionality of our neural network predictors), handcrafted gabor filters and GIST (spatial envelope) descriptors [72].

## 3 Results

### 3.1 How do trained models compare to randomly initialized models?

Previous work in the deep neural network modeling of mouse visual cortex found that a randomly initialized VGG16 predicted neural responses as well as, if not slightly better than, a VGG16 trained on ImageNet [9], suggesting that the neural predictivity of the features produced by a trained object recognition model are perhaps no better than the features produced by a randomly initialized one. Our results, on the other hand, suggest that the neural predictivity of trained versus randomly initialized models more generally depends on both the particular model being tested and the particular method used to produce the mappings between model and brain.

At the level of individual neurons (neural regression), 17 of the 91 model architectures we tested had randomly initialized variants that either matched or outperformed their ImageNet-trained counterparts. Replicating previous findings, we found these 17 architectures to include VGG16, as well as all 3 other VGG variants (11, 13 & 19), AlexNet, the DenseNet architectures (121, 169, 201), and almost all of the normalization-free architectures. Despite this, a paired t-test of the difference in scores across *all* models demonstrates that ImageNet-trained architectures are still overall more performant than their randomly initialized counterparts (Student’s *t* = 7.74, *p* = 1.37e — 11, Hedge’s 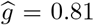). At the level of emergent representational similarity (RSA), ImageNet-trained models categorically outperform their randomly initialized counterparts, and by a large margin (Student’s *t* = 22.66, *p* = 5.81e – 39, Hedge’s 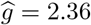).

Taken together, these results strongly affirm that *training matters*, and that randomly initialized features can only go so far in the prediction of meaningful neural variance. Differences between ImageNet-trained and randomly initialized models are shown in Figure 1.

**Figure 1:**
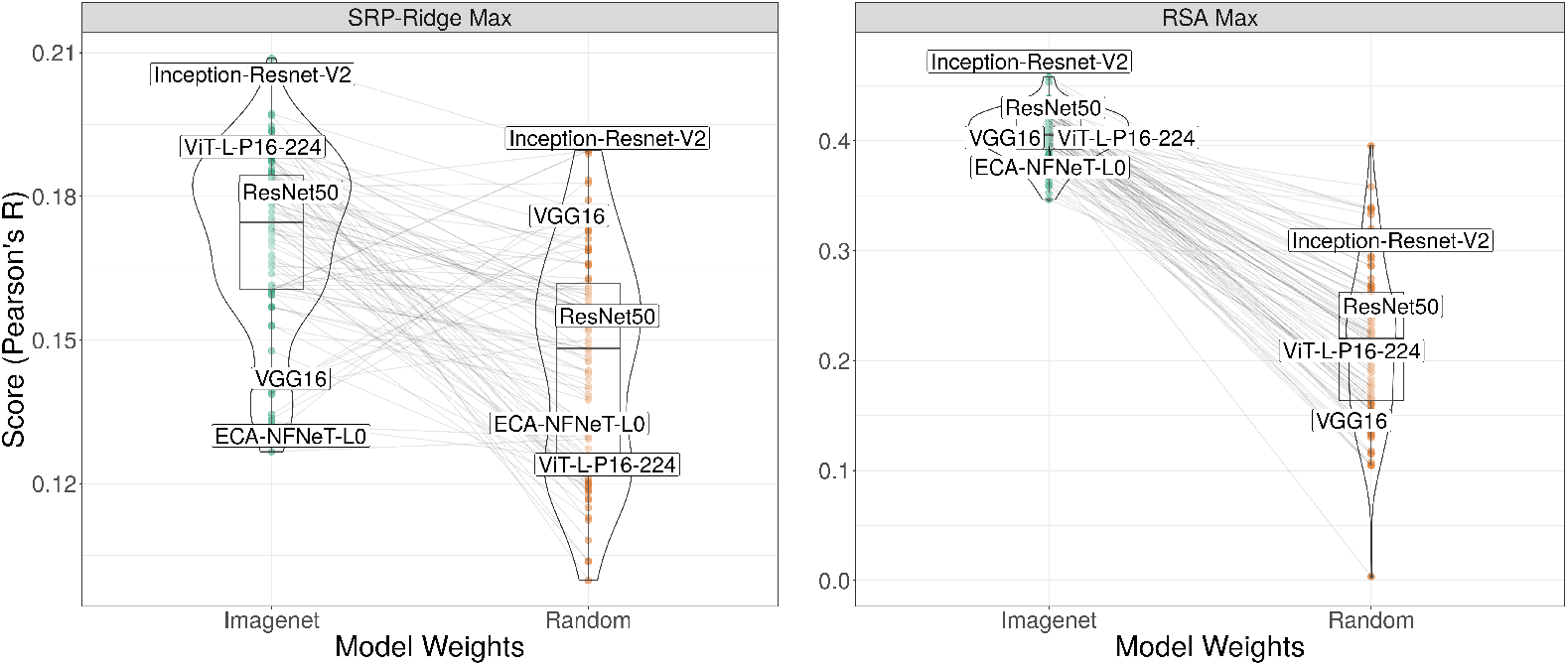
Each pair of connected points in these plots is the score of a trained versus randomly-initialized version of the same model architecture. Here, we’ve labeled 5 of these pairs for demonstration’s sake. The key takeaway of these plots is that ImageNet-trained models significantly outperform their randomly initialized counterparts – majoritively in the case of the SRP-Ridge Max metric, and categorically in the case of the RSA Max metric.

### 3.2 What kinds of architectures best predict rodent visual cortex?

The overall best architecture for predicting mouse visual cortex across both individual neurons (SRP-Ridge) and population-level representation (RSA) was an Inception-ResNet hybrid (Inception-ResNet-V2). There is a small, positive correlation between the depth of a model (the number of distinct layers) for both the RSA-Max metric and SRP-Ridge metric (Spearman’s *r* = 0.22, *p* = 0.001 and *r* = 0.192, *p* = 0.007, respectively), and a small, negative correlation for the total number of trainable parameters in the RSA Max metric (Spearman’s *r* = −0.18, *p* = 0.007). The latter of these is most likely driven by the relatively poor performance of parameter-dense architectures like VGG.

Markedly, trends previously noted in macaques [73] fail to materialize here. In particular, models with higher top-1 accuracies on ImageNet do not perform significantly better than models with lower top-1 accuracies. This relative parity is driven in large part it seems by newer models like EfficientNets, which across the board have dominant scores on ImageNet, but sometimes middling or poor scores in the predictions of rodent visual cortex we’ve tabulated here.

Compared to all other architectures, transformers on average fare slightly worse in the RSA Max metric (Student’s *t* = −3.96, *p* = 0.004, Hedge’s 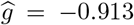), but moderately better in the SRP-Ridge Max metric (Student’s *t* = 2.45,p = 0.023, Hedge’s 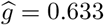).

Strikingly, transformers and MLP-Mixers boast the largest differences between ImageNet-trained and randomly initialized variants in the SRP-Ridge Max metric, with all pairwise t-tests significant at *alpha* = 0.05 after Bonferroni correction for multiple comparisons. This strongly suggests that the advantage of those randomly initialized variants that matched or outperformed their ImageNet-trained counterparts is an advantage conferred by properties of convolutional architectures (e.g., translation invariance), and not necessarily an advantage shared across random feature spaces writ large. (The rankings of these and other architectures may be found in Figure 8 in the Appendix).

**Figure 2:**
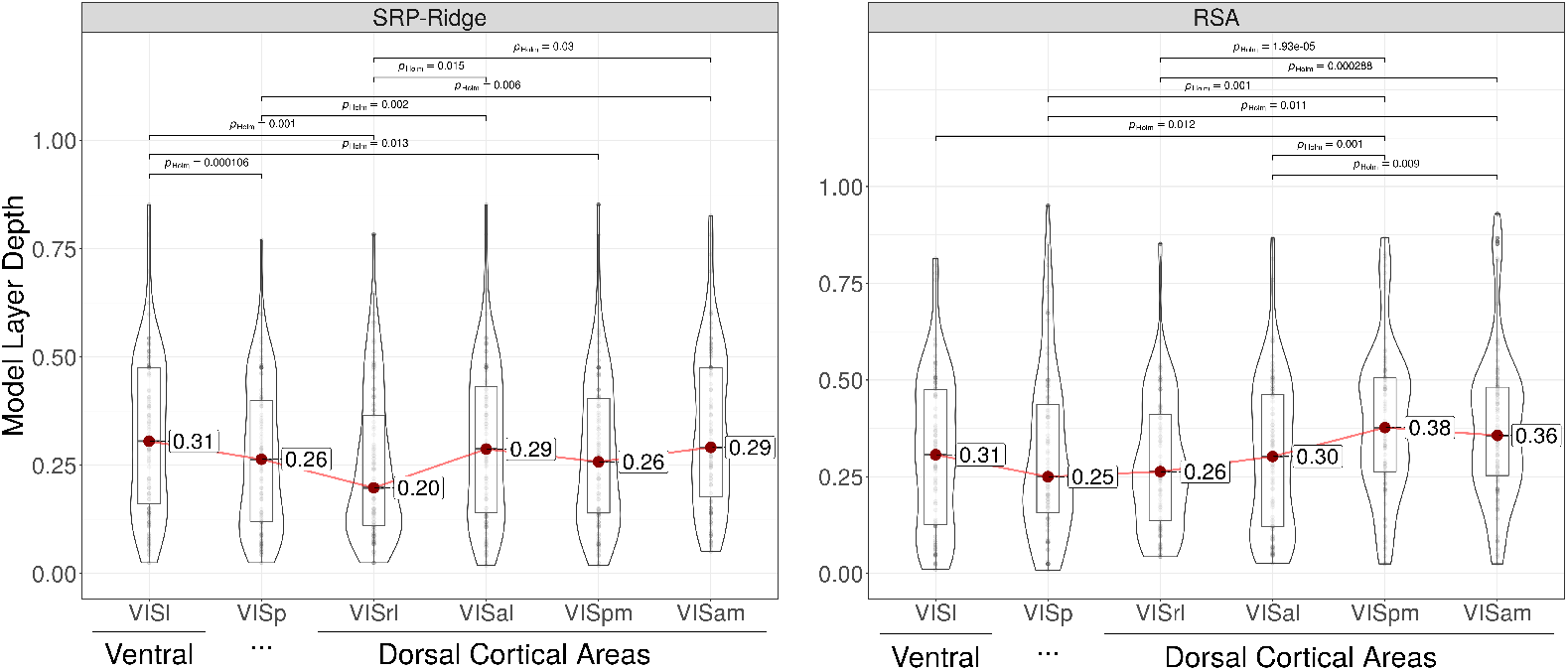
In these plots, we demonstrate a purely data-driven approach for recapitulating the proposed information processing hierarchy in mouse visual cortex. On the x axis in these plots are the 6 cortical areas in our optical physiology sample, arranged roughly according to the schematic proposed by [79] (Figure 3C). On the y axis is the average depth from 0 (the first layer) to 1 (the last layer) of the layer in each model that best predicts a given cortical area. Each point is an individual model. The horizontal brackets are drawn only in the case of a significant difference in a Games-Howell (Holm-corrected) pairwise comparison test. While noisy, these plots demonstrate that the deeper into the information processing hierarchy a given cortical area is, the deeper the average depth of the maximally correspondent model layers (and more complex the features). Even in those cases where we see a violation of the order suggested by the physiology (as in the case of the difference between in median depths between VISp and VISrl in the SRP-Ridge metric), these differences are insignificant. All *significant* differences point in the direction proposed by the physiology.

**Figure 3:**
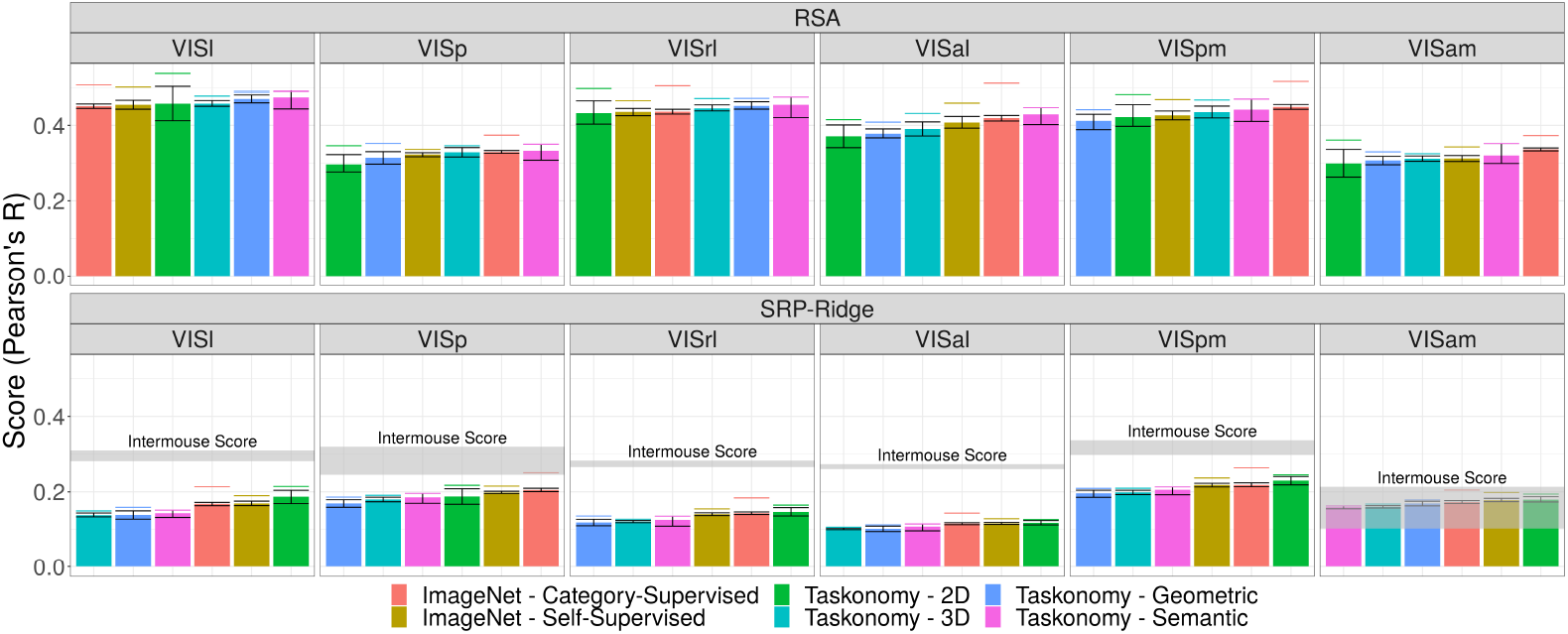
A mosaic of scores across distinct kinds of models in different cortical areas. Each bar is the average score across all models of a certain kind of training regimen or task affinity cluster (denoted by color). Error bars are 95% bootstrapped confidence intervals across models. The dashed lines above each bar are the scores of the best performing models in that category. Within each facet, the averages are arranged in ascending order, with the best performing models on the right and the worst performing on the left. For the SRP-Ridge metric, we’ve included a reference point we call the ‘intermouse score’, which leverages the neural activity of other mice to predict the activity in the cells of a target mouse. The band corresponding to the intermouse score is the 95% confidence interval across the scores for all cells in a given cortical area. Intermouse scores (as a sort of noise ceiling) tend to be a much more attainable target than the average splithalf reliability, which ranges from 0.886 to 0.906 across cortical area. There are a few takeaways from this plot: first, we see that there do not seem to be major ‘taskonomic’ dissociations across cortical area. 2D tasks (driven by unsupervised segmentation) dominate in the SRP-Ridge max metric, and despite a similarly strong showing from unsupervised segmentation, Semantic tasks are the evident superlative in the RSA Max metric. Second, we see that our models are in some cases on the verge of providing estimates of neural activity as well suited to the prediction of a given biological neuron as the reweighted activity of dozens of other biological neurons sampled from the same species. Finally, we see that (overall) there tend to be very few divergences or punctuated gaps in the average scores across model – underscoring that even sometimes *substantive* differences in the tools of computer vision do not always translate immediately to differences in neural predictivity.

**Figure 4:**
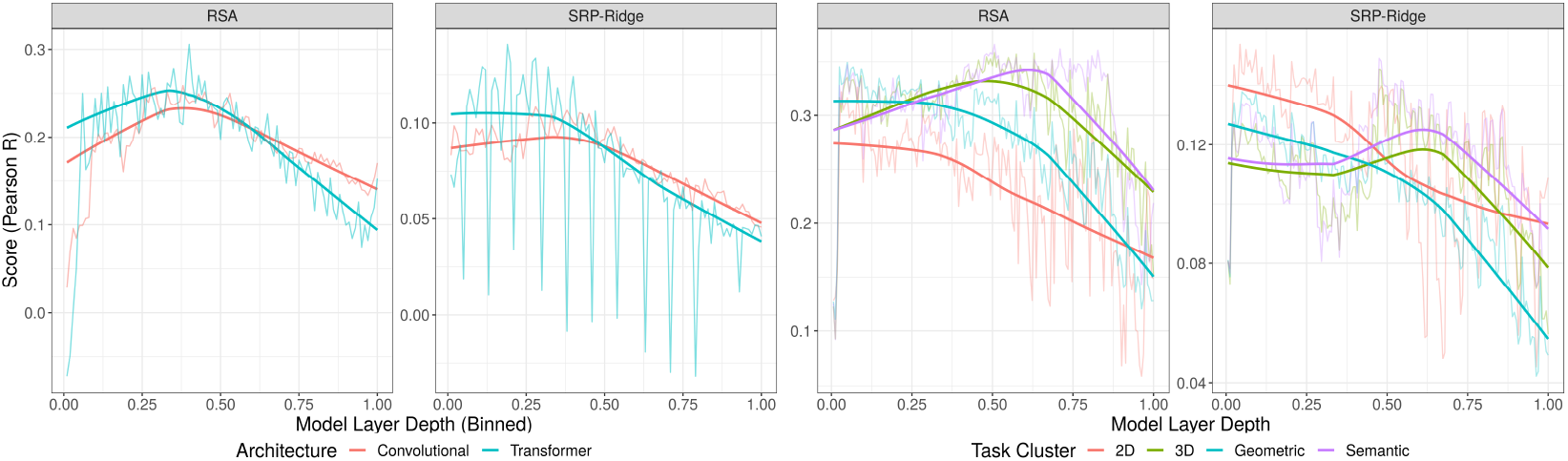
Aggregate layer by layer comparisons across ImageNet-trained models (left) and Taskonomy models (right). The horizontal axis shows relative model layer depth, from the first to the last layer. Because the models on the left vary significantly in size, we discretize the relative depths into bins of size 0.01 (100 bins). The jagged, semitransparent lines are the mean scores at a given depth. (These are particularly jagged for transformers due to the heterogeneous nature of the computational blocks, which engenders large troughs in predictive power). The opaque, smoothed trend lines are the outputs of locally weighted linear regressions (lowess) computed at each point with a span of 2/3 the width of the axis. Plots show that convolutional networks and transformers have similar predictive power at similar depths despite their significant computational differences. (right) Features most predictive of cortical responses across the Taskonomy models vary in their relative depth. The reversal in the ranks of 2D models between RSA Max and SRP Ridge Max is seen as a vertical shift in the intercept of the curve relating depth to score – though note that the slope and shape of the curve remain similar across metrics. 2D models are most predictive in the early layers, but are nevertheless superseded by other tasks in the later layers at the level of representational similarity.

**Figure 5:**
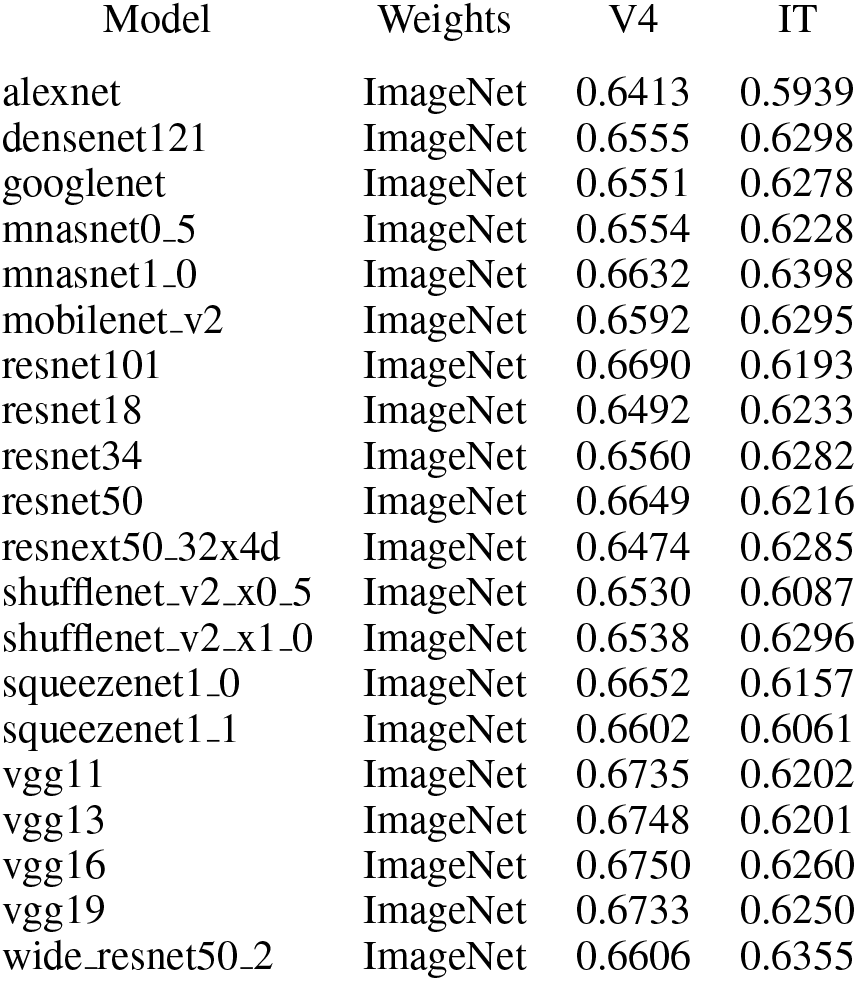
Scores for a subset of models tested on the macaque V4 & IT benchmarks of BrainScore. Corresponding scores for models in our set that overlap with models tested by the BrainScore team may be found at: https://www.brain-score.org/

**Figure 6:**
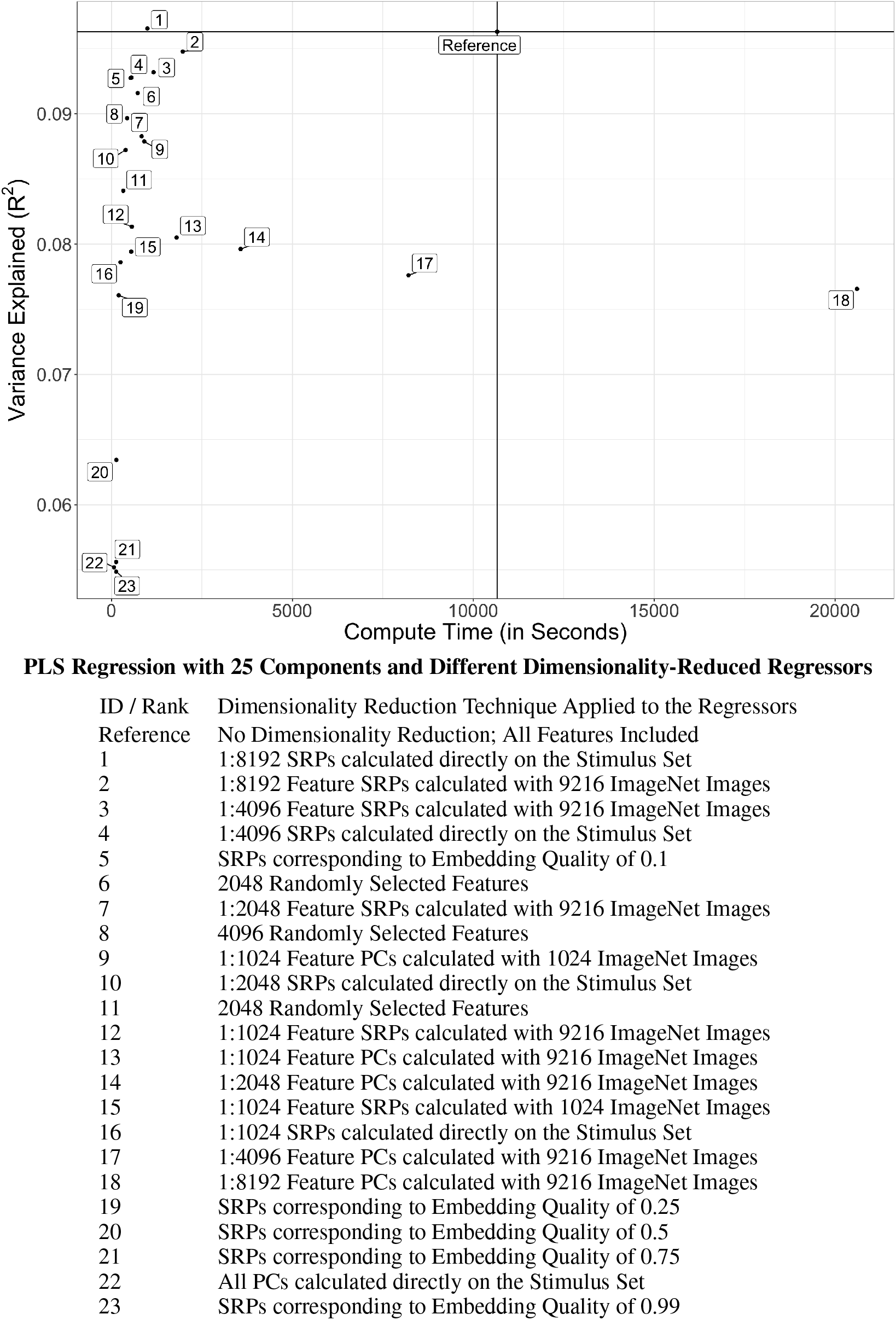
Variance explained versus compute time for a PLS Regression with 25 components and different types of dimensionality reduction techniques applied to the regressors. Descriptions of the dimensionality reduction steps associated with each data point are provided in the table above. The vertical and horizontal ablines triangulate a reference point for all methods: that is, a regression in which all features from a given model layer are used simultaneously without dimensionality reduction. Notice, that at least one PCA-based method (18) takes longer than this full regression.

**Figure 7:**
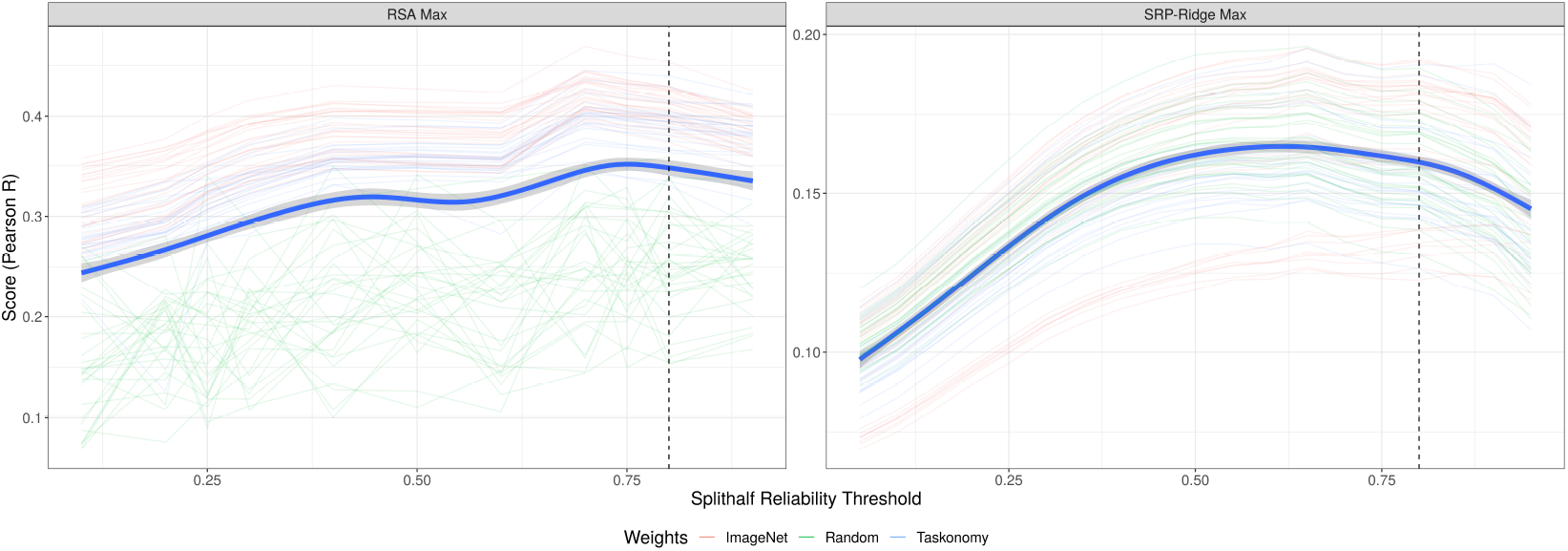
Scores for a variety of models with both the RSA Max (left) and SRP-Ridge Max (right) metrics at different levels of reliability thresholding. The jagged, semitransparent lines are the scores for individual models. The smooth, opaque line is the output of a generalized additive smoother fit across all models. (Error bars are bootstrapped 95% confidence interval across models). The dotted vertical line is the threshold we use in the main analysis. Based on these results, one might argue a more performative threshold would have been closer to 0.7 or 0.75. In the future, we plan to more closely emulate methods designed to derive the optimal threshold empirically (e.g. reliability-based voxel selection in human fMRI [104]).

**Figure 8:**
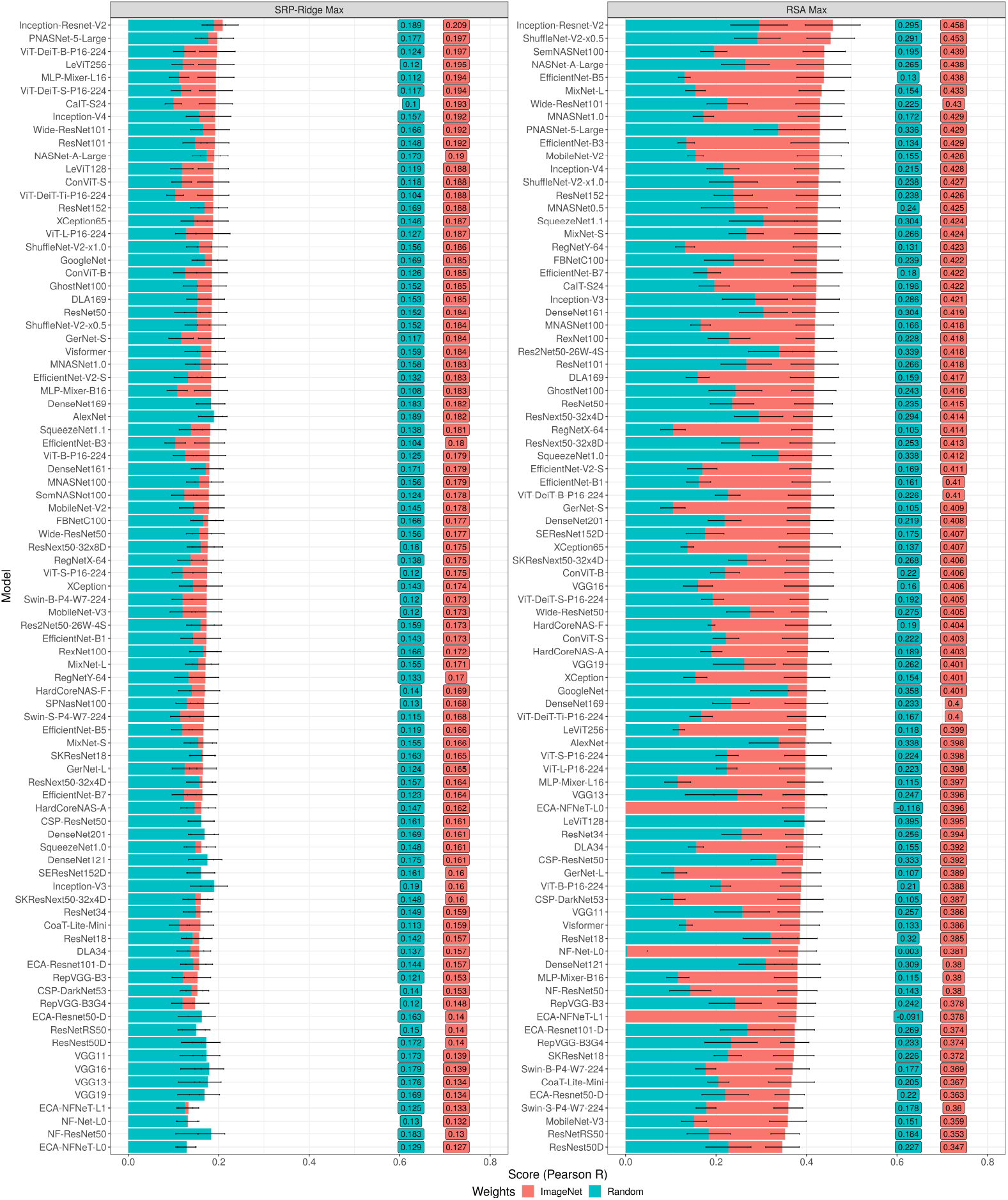
Rankings across model architectures, sorted by the scores of the ImageNet-trained variants (red) of each. Instances in which the randomly initialized variants (blue) outperform their ImageNet-trained counterparts are visible in those rows where the blue entirely overlaps the red. Error bars are 95% bootstrapped confidence intervals across the 6 cortical areas.

### 3.3 What kinds of tasks best predict rodent visual cortex?

The overall best Taskonomy encoder across both the RSA and SRP-Ridge Max is 2D segmentation (ranking second and first respectively; see Figure 9 in the Appendix). At the level of individual neurons (SRP-Ridge), 2D tasks (keypoints, autoencoding, inpainting) dominate. At the level of representational similarity (RSA), all 2D tasks but 2D segmentation fall to the bottom of the rankings, and Semantic tasks (object recognition and semantic segmentation) rise to 2nd and 3rd place.

**Figure 9:**
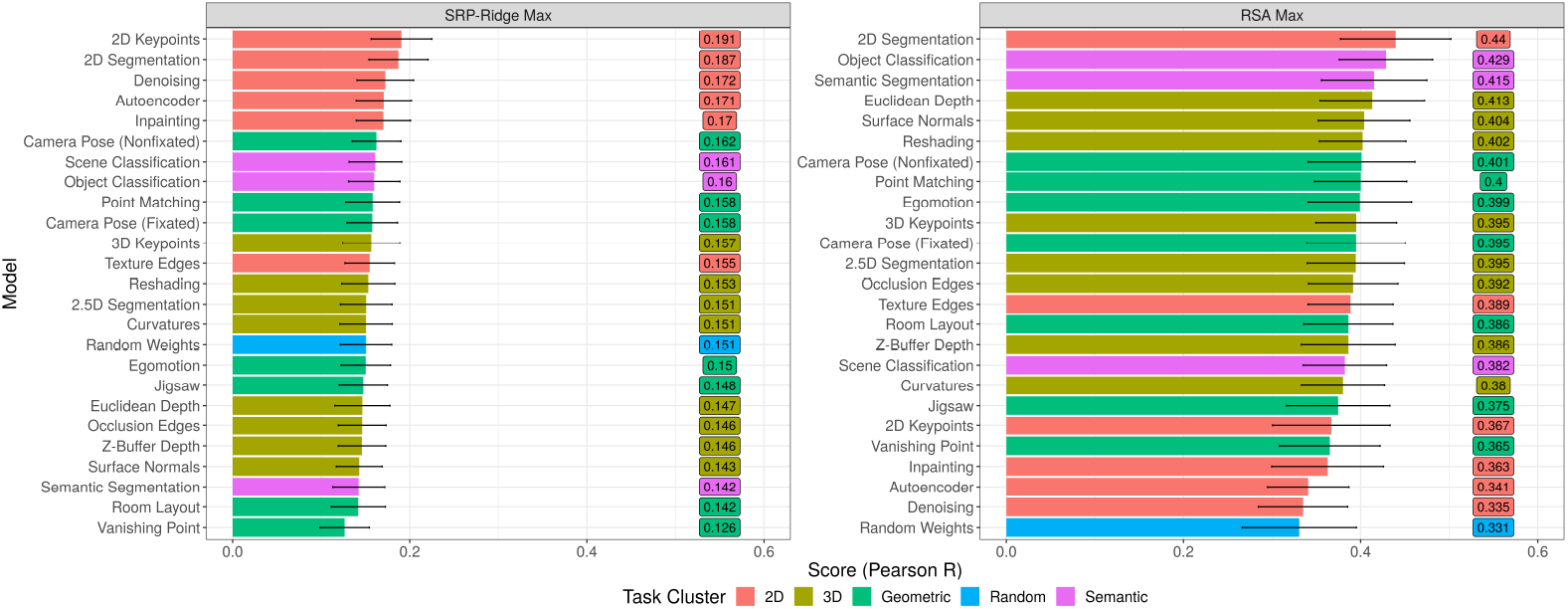
Rankings across Taskonomy encoders, color coded by the Taskonomic cluster to which each belongs. Notice the contrast between the SRP-Ridge Max (which favors 2D tasks) and RSA Max metric (which favors Semantic tasks), but also the relative rank of 2D Segmentation in both.

This reshifting in rank presents a curious case for interpretation, suggesting most likely that while the representations of individual neurons may be coordinated more by the lower level, less abstract features necessary for performing well on most 2D tasks, the overall neural population codes are coordinated more by the parsing of the visual input into ethologically and spatially relevant units via the segmentation and classification tasks. Notably, the original research from which these PyTorch models were adopted offers an auxiliary data point that may anchor this interpretation more concretely. The top 3 models in our RSA Max metric (2D segmentation, object classification, semantic segmentation) are likewise in the top 5 of a ranking the original researchers produced by pitting the Taskonomy encoders against one another as pretrained ‘perceptual systems’ for reinforcement learning agents learning to navigate a virtual environment (see [54], figure 13 in the appendix). This raises the possibility that the reason these models are optimally predicting the visual neural population code for mice is simply because that code is coordinated in service of navigation.

**Figure 10:**
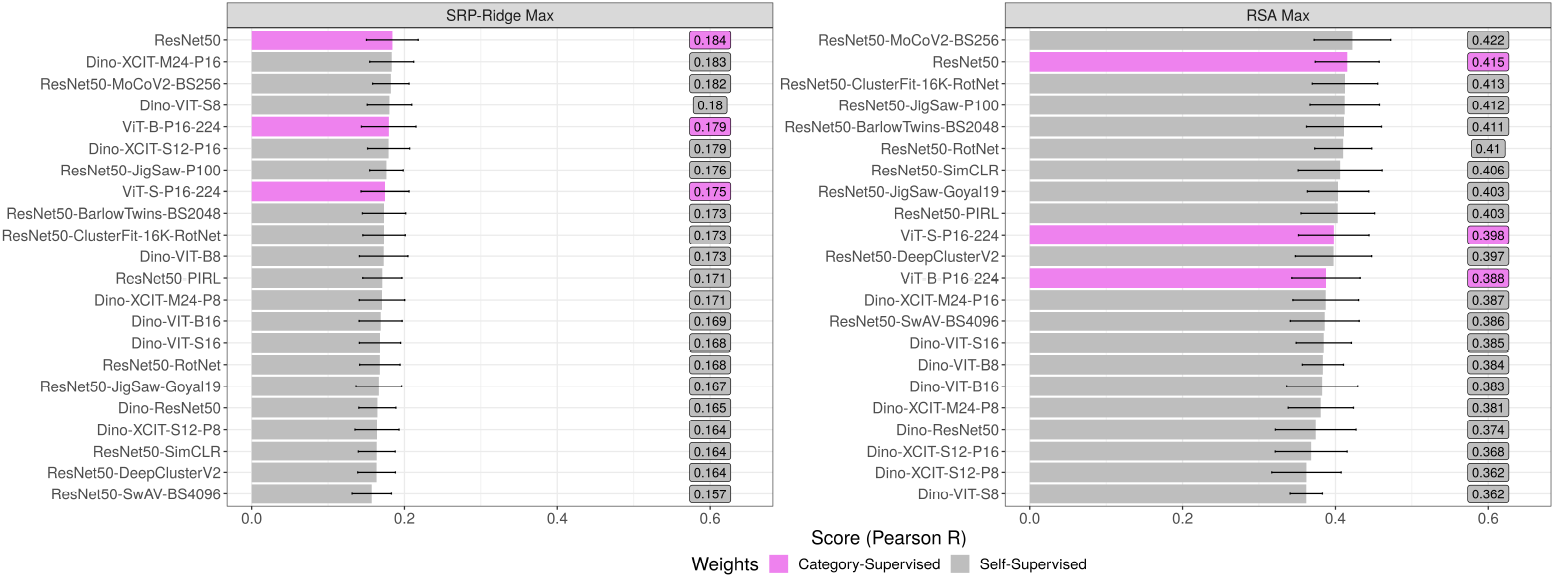
Rankings across self-supervised models. Where available, the category-supervised version of a given model architecture is shown in violet.

**Figure 11:**
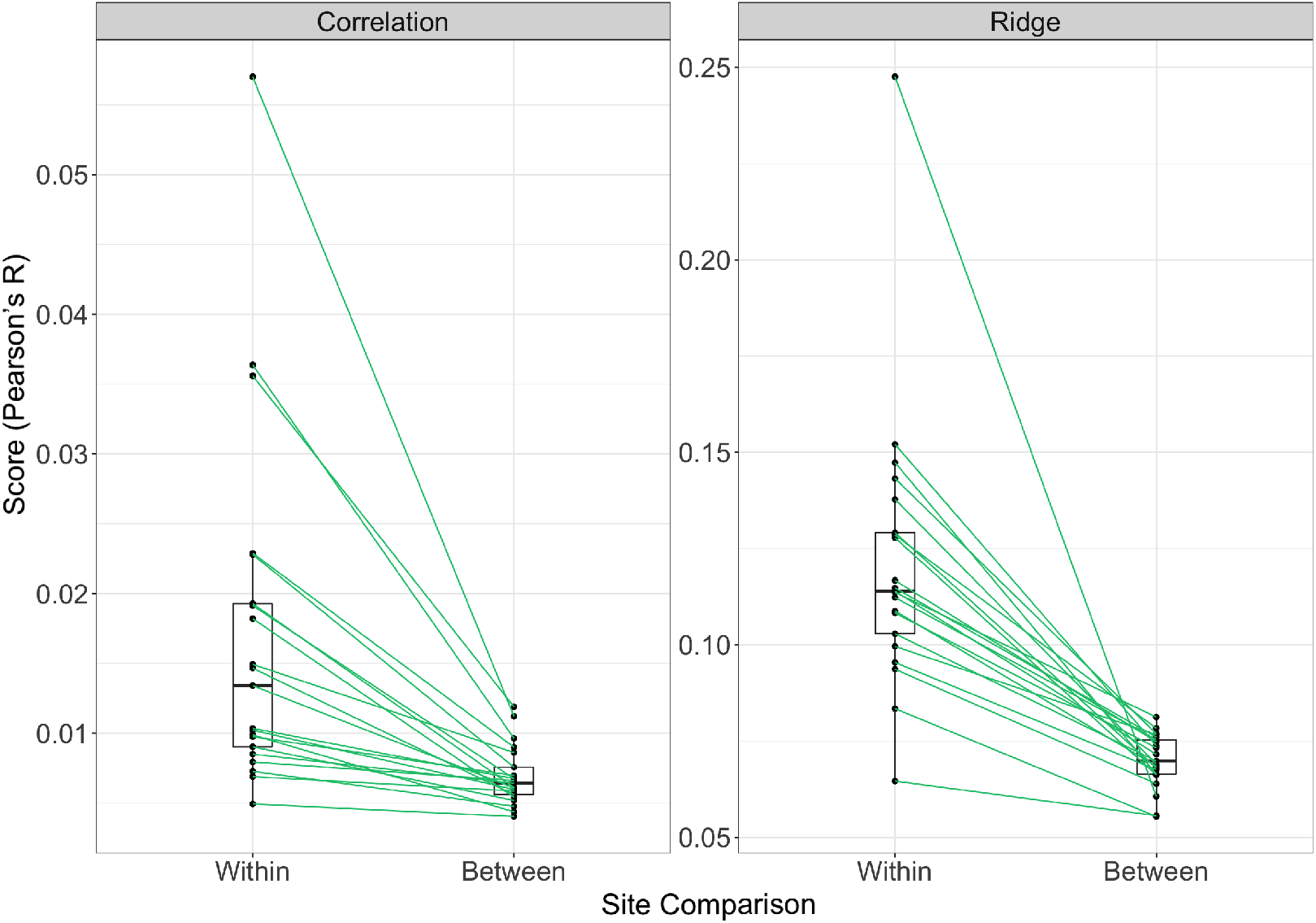
Differences in the predictive power of neurons predicting other neurons *within* site versus neurons predicting other neurons between site. Each paired set of points in this plot is a distinct cortical site (cortical area + cortical layer – 21 in total). The Correlation metric is the average of up to 1000 pairwise comparisons between cells from within the same cite and cells from other sites. The Ridge metric is the iterative prediction of individual neurons using up to 1000 neurons from within the site versus 1000 neurons from other sites. (The number of predictors was subsampled to ensure an equal number of neurons contributed to the within versus between samples). What this plot demonstrates is that neuroanatomical regions do in this case meaningfully correspond to regions with distinct representational profiles, a fundamental component of functional specialization.)

**Figure 12:**
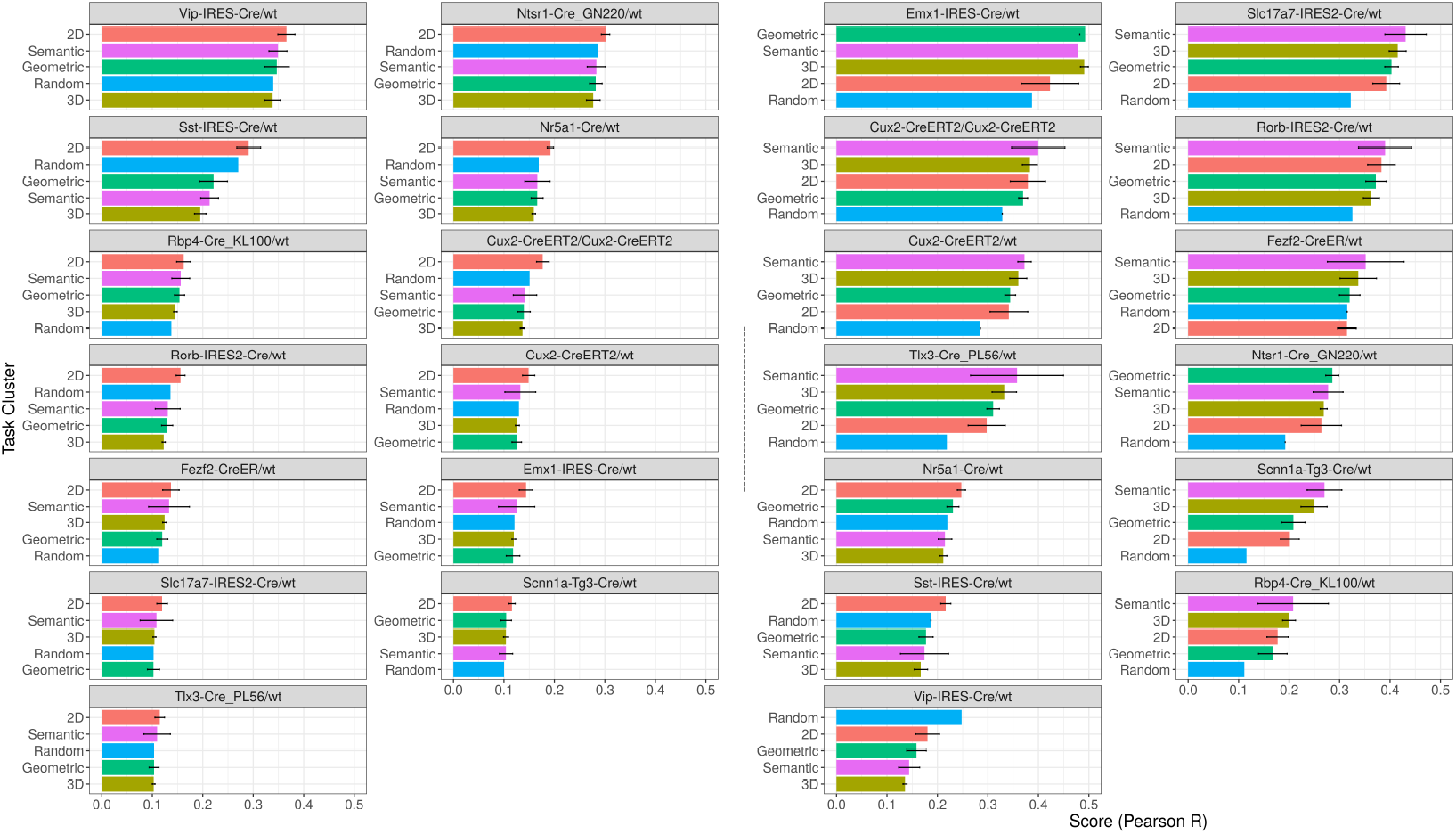
Taskonomy scores across genetic cre line. On the left is SRP-Ridge Max metric; on the right is the RSA Max metric. The facets are shown in descending order by the overall magnitude of their mean scores. Of note: Large-scale motifs present when aggregating across cortical area (such as the dominance of 2D models in SRP-Ridge; the dominance of semantic models in the RSA Max; and the lack of clear Taskonomic dissociations) are recapitulated across cre line. Nevertheless, some differences are salient. For example, the aggregate scores for certain cre lines using the SRP-Ridge Max metric are much higher on average than the scores obtained in any cortical area.

**Figure 13:**
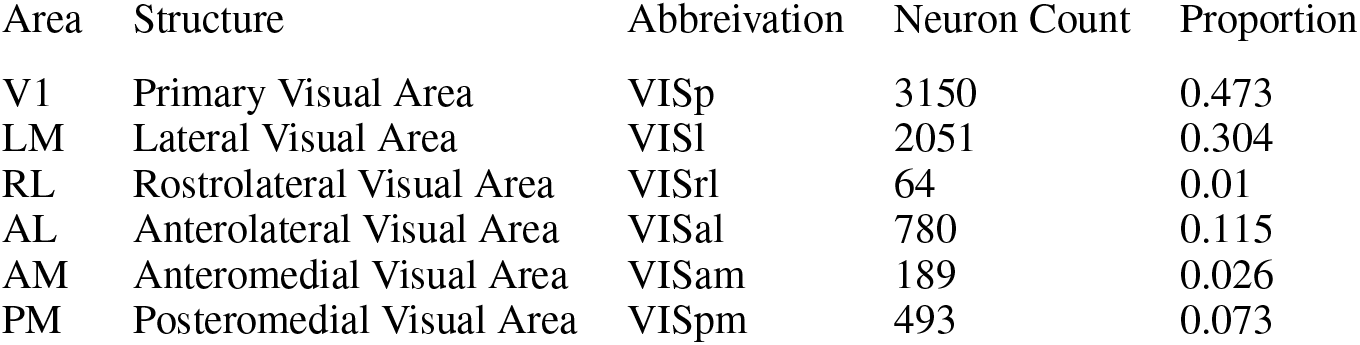
A glossary of areas in the mouse visual cortex.

### 3.4 How do category-supervised models compare to self-supervised models?

Whether with ResNet50 as their base, or a vision transformer, self-supervised models seem to be verging closer and closer to the predictive power of their category-supervised counterparts. Our most predictive self-supervised ResNet50, for example, (a MocoV2 model) effectively matches its category-supervised counterpart in the SRP-Ridge Max metric (with scores of .182 and .184, respectively), while slightly outperforming its category-supervised counterpart in the RSA Max metric (with scores of .422 and .415, respectively). While this single comparison by no means denotes a statistically significant superiority of self-supervised models (which would require training multiple iterations of each), it does begin to provide preliminary evidence for parity.

### 3.5 How well do non-neural network baselines predict rodent visual cortex?

Non-neural network baselines somewhat uniformly fail to predict neural activity as accurately as deep net features (though see Section A.10 of the Appendix for a counterexample). We tested three baselines: 1) a bank of Gabor filters of applied to 8×8 grids of each image; 2) the PCs of the resultant feature matrices (i.e. the Gist descriptors [72]); 3) and the max across 600 iterations of 4096 random Fourier features (a dimensionality matching that of our SRPs). Ridge regressed with generalized cross-validation, these feature models yield average scores of 0.07, 0.06 and −0.014, respectively. Compared via representational similarity, they yield average scores of 0.20, 0.25 and 0.011.

### 3.6 How ‘deep’ are the layers that best predict rodent visual cortex?

Echoing previous results [7, 8], we find across all ImageNet-trained architectures, regardless of metric, that the features most predictive of rodent visual cortex are found about a third of the way into the model (though see Section A.5 of the Appendix for some caveats). These early to intermediate visual features go beyond basic edge detection but are far from the highly abstracted representations adjacent to final fully connected layers. Across Taskonomy encoders, 2D & Geometric tasks yield their best features in earlier layers; 3D & Semantic tasks yield their best features in more intermediate and later layers. Note that these aggregate motifs do not preclude subtler differences across cortical area, which we discuss in the section below.

### 3.7 Are there differences in model predictions across cortical area?

In this work, we address this question from two perspectives: that of hierarchy and that of function. In primate visual cortex, it is common consensus that there exists a distinct information processing hierarchy along the ventral visual stream [74–76], with posterior sites like V1 and V3 defined by features like oriented edge detectors, and more anterior sites like V4 and IT defined by more complex morphologies. While there continues to be some debate as to whether a similar hierarchy exists in rodent visual cortex, a large body of anatomical, functional and physiological work [77–83] has coalesced around a meaningful hierarchy that consists first of a ventral / dorsal split after primary visual cortex (VISp), with VISp leading to VISl in the ventral stream and VISp leading to VISrl - VISal - VISpm - VISam in the dorsal stream. Strikingly, our modeling does seem to provide corresponding evidence for this circuit in the form of a data-driven hierarchy produced purely by taking the median depths of the model layers that best predict the neural activity in each of these cortical areas, and assessing for difference across them. A nonparametric ANOVA shows an overall difference in depth across cortical area to be significant for both our SRP-Ridge metric (Friedman’s *χ*^2^ = 34.08, *p* = 2.29e – 06, Kendall’s 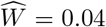) and our RSA metric (Friedman’s *χ*^2^ = 37.05, 5.86e – 076, Kendall’s 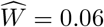). Subsequent pairwise comparisons show many of the differences that underlie this group-level effect to be differences between earlier and later layers of the information processing hierarchy established in the literature. (For further details, see Figure 2.)

Other differences across cortical area that we might expect are differences driven by function. Research into primate visual cortex over the last two decades has unveiled a significant degree of functional organization over and above purely anatomical organization [84–86], with distinct subregions defined in large part by their differential activity in response to different kinds of stimuli. To try and replicate this in mouse visual cortex we search for Taskonomic organization, a proxy of functional organization wherein distinct neural sites are better or worse predicted by the features from different taskonomy encoders. Curiously, and in contrast to previous findings in human fMRI [55], it seems to be the case that the scores of different Taskonomic clusters are relatively consistent across cortical area; see Figure 3.) This suggests that mouse visual cortex *may* be more functionally (or Taskonomically) homogenous than primate visual cortex, with anatomical descriptors providing little to no cue of functional difference – though this seems unlikely given other analyses we’ve performed showing greater similarities of neurons *within* cortical site than *between* cortical site (see Section A.11 for details). Another (more likely) alternative is that the tasks of computer vision are just not so neatly mapped onto the tasks of biological vision in mice.

### 3.8 How do the predictions compare across RSA and neural regression?

While prior work has addressed this question theoretically [87], it’s rarely the case that representational similarity and neural regression are compared directly and empirically. Here, we compare our RSA and SRP-Ridge metric both at the level of overall rankings (taking the max across layers) and at the level of individual layers, the latter of which provides a much more detailed assessment of how different feature spaces map to cortical representation.

In terms of overall rankings, the Spearman rank order correlation between the two methods is either 0.56 (*p* = 8.36 × 10^-19^) or 0.59 (*p* = 3.17 × 10^-12^), depending on whether you include or exclude the randomly initialized architectures. In terms of layer by layer comparisons, we decompose the Spearman correlation across distinct combinations of model and cortical area. The average coefficient between the two methods, along with bootstrapped 95% confidence intervals is 0.468 [0.447,0.489] or 0.626 [0.613,0.639], again depending on the inclusion or exclusion of the random models. This suggests a significant degree of overlap between the kinds of features that optimally predict the representations of both individual neurons and neural populations. Of course, the averages here do obscure some meaningful subtrends and idiosyncrasies. For details, see Figure 4.

### 3.9 How well are we doing overall in predicting mouse visual cortex?

The overall best model in any cortical area across either of our metrics is unsupervised 2D segmentation in anterolateral visual area (VISal), with an RSA Max score of 0.538. The (Spearman-Brown) splithalf reliability of the RDM for this area (an effective proxy of its explainable variance) is 0.89. This means our most predictive model in any cortical area across any metric is little more than halfway to the noise ceiling.

Of course, it’s possible this noise ceiling is a bit too strict. Instead of requiring the model to predict the neural data as well as the neural data predicts itself, another possible target to which we might recalibrate is the relative performance we would expect if (instead of an artificial neural network) we used the responses of another biological network as the model to predict neural activity. Inspired by recent work [88], and to better contextualize the scores of our SRP-Ridge metric, we attempted a version of this here. To compute this reference, we proceeded again neuron by neuron using the exact same neural regression method (dimesnonality reduction, and hyperparameters) described in Section 2.5.2, but instead of using the responses of a deep net layer as the predictors in our ridge regression, we used the responses of the neurons from the same cortical area in all other mice (conspecifics) across the donor sample. Conceptually, this ‘intermouse score’ represents how well we might do if our model of a given mouse brain were other mouse brains.

Averaging across both cortical area and model, the average distance (with 95% bootstrapped confidence intervals) between the best performing deep net feature spaces and the mean of the intermouse scores (expressed in the same units of Pearson’s *r* we’ve used heretofore) is 0.0985 [0.0940, 0.103]. Compare this to the same distance computed relative to the splithalf reliability: 728 [0.726, 0.731]. On average, then, while our artificial models are capturing only a fraction of the total explainable variance relative to the splithalf noise ceiling, they’re verging increasingly close to the predictive threshold suggested by the reweighting of biological neurons from the same species. The performance of models relative to the intermouse score may be seen in the lower half of Figure 3.

## 4 Discussion

Our intent with this work was to provide a preliminary atlas for future ventures into the deep neural network modeling of rodent visual cortex. To this end, we have deliberately invested in introspective analyses of the tools we used, as well as the curation of deep neural networks we hope will provide informative waypoints. Obviously, the atlas is far from complete. Other model classes like recurrent models [89, 90], equivariant models [91], and robotic models (e.g. for visual odometry [92]) are promising candidates for inclusion in future benchmarks, and our neural encoding & representational similarity metrics are just two of many variants.

Nevertheless, the results we have presented here indicate that neural recordings from the visual brains of mice can (with care and caution) be compared to deep neural networks using many of the same tools we’ve used to better characterize the visual brains of monkeys. Having as reference two animal models that occupy very different ecological niches and are separated by tens of million years of evolution makes it far more likely that insights into vision gleaned across both are actually fundamental to perceptual meaning-making and not just some idiosyncratic quirk specific to any one evolutionary trajectory. Primate and rodent vision do differ rather drastically, even in fairly basic ways: mice lack a fovea, have a retina dominated by rods for vision under low light, and spatial acuity less than 20/1000 [93], making their primary visual system more akin to the primate peripheral system – and making it all the more curious that the same models explain decent amounts of variance in both. The differences between the species, it seems, may not be so irreconcilable at the level of modeling, but only with future work more carefully controlling for distinct aspects of each organism’s unique physiology (see Section A.5) can more concrete conclusions of this kind be made.

Beyond considerations of distinctive physiology is the indispensable point that perceptual systems should always be considered in service of behavior. It’s possible that mice mostly rely on vision as a sort of broad bandpass filter for lower-frequency, dynamic stimuli the animal can then flee, fight, or further investigate with its whiskers — perhaps its most sophisticated sensory organ [94]. Another possibility is that mice use vision to facilitate navigation. The dominance in our Taskonomy results of 2D segmentation, object recognition and semantic segmentation (all tasks that have elsewhere been shown to provide effective, transferable features for the simulation of robotic navigation) provide some evidence for this. Of course, the behavioral roles of rodent vision may very well be manifold. Understanding this plurality in a readily available model species could in the end be key for bridging the gaps that remain between biological and computer vision [95]. The unparalleled access, resolution, and control afforded by rodent neuroimaging have already revolutionized our understanding of the relationship between perceptual representation and behavioral output. Combined with novel methods like the embedding of neural networks in virtual agents [96] in ecologically realistic environments, this kind of data may well provide a testbed for better situating the tasks of computer vision in the broader behavioral context of agentic scene understanding.

In summary, only novel combinations of architecture, task and mapping will help to explain the highly reliable neural variance we’ve yet to explain in our current survey. Already this recombination is under way: Shi et al. [97] have created a custom CNN designed specifically to match (processing stage by processing stage) the anatomy of rodent visual cortex, while Nayebi et al. [88] have combined the power of self-supervised learning with smaller, shallower architectures to more fully account for the ethological realities of rodent behavior and the differences in computational bandwidth that shape and constrain their visual systems. More work of this variety will be necessary to more fully model the rich diversity and fiendish complexity of biological brains at scale – even the very smallest ones.

## 4.1 Acknowledgements

We thank Martin Schrimpf, Tiago Marques, Jim DiCarlo, as well as many others on the BrainScore team for helpful discussion, feedback, and inspiration. We would also like to thank the Allen Institute founder, Paul G. Allen, for his vision, encouragement, and support.

## 4.2 Code Availability

More results and code for the replication of our analysis may be found at this GitHub repository: github.com/ColinConwell/DeepMouseTrap (License GPL v2)

## 4.3 Compute Required

We used a single machine with 8 Nvidia RTX 3090 GPUs, 755gb of RAM, and 96 CPUs. GPUs were used only for extracting model activations, and could (without major slowdown) be removed from the analytic pipeline. Dimensionality reduction and regression computations were CPU and RAM intensive. Replicating all of our results would take approximately two weeks on a similar machine.

## 4.4 Ethics Statement

Lest our science forget the life that powers it, we must note that behind the phenomenal dataset provided by the Allen Institute are 256 laboratory mice, each of which was subjected to multiple surgeries, a highly invasive neuroimaging technique and genetic engineering. The moral parameters of this particular praxis of neuroscience are contentious, and not without reason. While we believe centralized, comprehensive and (most importantly) public datasets like those provided by the Allen Institute may actually decrease the total number of laboratory animals required for similar kinds of empirical projects, we acknowledge with solemnity the cost to life required.

## A Appendix for Neural Regression, Representational Similarity, Model Zoology & Neural Taskonomy at Scale in Rodent Visual Cortex

### A.1 Does our neural regression method work?

To ensure our neural regression method works, we verify its efficacy on a known benchmark: the activity of 256 cells in the V4 and IT regions of two Rhesus macaque monkeys, a core component of BrainScore [4]. BrainScore’s in-house method involves a combination of principal components analysis (for dimensionality reduction) and k-fold cross-validated partial least squares regression (for the linear mapping of model to brain activity). Here, we exchange principal components analysis for sparse random projection and partial least squares regression for ridge regression with generalized cross-validation. We compute the scores for each benchmark in the same fashion as BrainScore: as the Pearson correlation coefficient between the actual and predicted (cross-validated) activity of the biological neurons in the V4 and IT samples.

Taking for example a standard AlexNet architecture, our neural regression method yields gains of 16.5% (from 0.550 to 0.641) & 16.9% (from 0.508 to 0.593) on reported scores for V4 and IT, respectively. Across 6 other Torchvision architectures we tested with scores posted on the BrainScore leaderboard, our method yields gains on average of 13% for V4 and 23% for IT, and at its best yields a gain of 34% for SqueezeNet1-0 predicting IT. Though these scores are provisional (since official BrainScore results involve an additional step of validation – ‘commitments’ – on data not publicly available), we consider this a strong validation of our neural regression method, which is both less computationally expensive, far faster and (to the extent that the generalized cross validation represents an optimal approximation of how well the mappings fit to our models might generalize to novel biological samples) more accurate than the combination of PCA and PLS. (Speed tests may be found in Section A.3 of this Appendix.) Figure 5 shows the scores for the full subset of models we tested on primate BrainScore.

### A.2 Deeper Dive: How do our neural benchmarking methods compare to others?

In the main analysis, we roughly group existing methods for comparing the responses of deep neural networks to neural responses recorded from brain tissue into two categories: neural regression and representational similarity analysis. In reality, this division is often not so neatly dichotomous. Some of the first brain-to-network comparisons availed themselves of both these methods simultaneously; Yamins et al. [1] citing Carandini et al. [68] and Kriegeskorte et al. [66] used linear regression for mapping responses in individual neural sites and representational similarity analysis for populations. Other seminal work comparing deep nets to primate visual cortex pioneered distinctive variants of each. Güçlü and van Gerven [67] employed regression in the form of encoding models to assess the hierarchical correspondence between earlier and later layers of processing in vivo and silico. Khaligh-Razavi and Kriegeskorte [98] built representational dissimilarity matrices by “remixing” and “reweighting” model features according to their performance in a support vector machine classifier trained on major categorical divisions in the stimulus set. Zhuang et al. [58] citing Klindt et al. [99] uses a form of masked regression to better account for spatial information (e.g. properties of the receptive field) in the target feature spaces. In the context specifically of comparisons to rodent neurophysiology, Cadena et al. [9]’s neural encoding method predicts spike rate with a core feature model (VGG16) in tandem with a “shifter” network and “modular” network that correct for extraneous influences on recorded brain activity (including eye movements and running speed). A possible third strain of methods that doesn’t fit so neatly into the binary of regression versus representational similarity are canonical correlation and alignment methods. These techniques leverage what is often assumed to be an underlying latent space of similarity shared across divergent high-dimensional datasets to assess (via projection) the shared variance between them. Canonical correlation and alignment methods are popular in both the machine learning [100, 101] and neuroimaging communities [102], but have so far been applied mostly to comparison within, rather than across, domains and neural substrates. The relative advantages of these various approaches as they pertain to characterizing the representational structure of biological brains is largely uncertain, with a comprehensive comparison of techniques on the same dataset seemingly absent from the literature.

The current standard for high-throughput benchmarking of neural data on neural models is perhaps that of Schrimpf et al. [44] in BrainScore, a method that consists of a partial least squares (PLS) regression fit individually to each neural site (in their case, a cluster of neurons around a given electrode in a microarray), wherein the regressand is the responses from that site and the regressors are the principal components of a target model’s feature space. The end product of this process is a Pearson correlation coefficient (unadjusted or reliability-corrected) quantifying the relationship between actual neural activity and the neural activity predicted by the linear mapping. While effective, this combination of principal components analysis and partial least squares regression tends to be a computationally expensive process – often prohibitively so in the absence of cloud or cluster computing. The final approach we use in the primary manuscript is a more computationally efficient version of this process. The reasoning behind the particular neural regression we use (assessing the tradeoff between accuracy and computational traction) may be found in the section below.

### A.3 How do different neural regression methods trade off in terms of speed & accuracy?

Given the many variants of neural regression used in the analysis of the human and nonhuman primate brain (and to a lesser extent the rodent brain), we experimented with a number of possible approaches before settling on the one detailed in the primary manuscript. Attempting to directly mirror the approach described in Schrimpf et al. [4], we began with a method combining principal components analysis for dimensionality reduction with partial least squares regression for neural prediction. So as to capture more dimensions of variance in a given model’s feature spaces, and not ‘double dip’ meaningful dimensions of variance with the regression to follow, Brain-Score computes a set of principal components on the features from an auxiliary set of held-out ImageNet images, then extracts the loadings of the features from the target stimulus set on these same components. These loadings are subsequently made the regressors in a partial least squares regression of 25 components, with a given neural site (the activity from a microelectrode array) as the regressand. The most computationally intensive step of this process is the calculation of the PCA on the features from the auxiliary Imagenet images – requiring in larger models like VGG16 upwards of 450GB of RAM for a single layer. The prohibitively large expense of this PCA prompted us to search both for alternative dimensionality reduction techniques, as well as for the possibility of extracting fewer total dimensions with whatever technique we chose. A summary of the outputs of this search (using only a small subset of models and a fraction of our total neural data) may be found in Figure 6.

What this search made clear (at least in the context of our specific neural data and stimulus set) is that approaches involving PCA were doubly suboptimal, taking orders of magnitude longer to compute, and actually costing a nontrivial portion of score. In the PLS regressions as well, it quickly became clear that 10 components yielded scores comparable (if not equivalent) with those of twice or thrice as many components, suggesting nothing was to be gained from more components apart from longer compute times. Finally, while somewhat less definitive than the 2 previous points, our tests did suggest that using 4096 sparse random projections was roughly comparable to using 8192 sparse random projections – and translated to about 0.75 of the compute time per projection. When in an additional test a ridge regression performed in roughly 0.29 seconds what a PLS regression with 10 components performed in roughly 1.31 seconds (representing an over 200% percent gain in speed, and consistent across 10 iterations), while producing only a 0.004 difference in average R^2^, we abandoned PLS regression entirely in favor of ridge regression. Switching to ridge regression meant we could also make use of generalized cross-validation [70, 103], cutting the time required for k-fold cross-validation from roughly 1.05 seconds to 0.25 seconds. While it is most certainly the case that these tests were not comprehensive enough in terms of models or neural data to cover the full range of contingencies and idiosyncrasies of analysis, we felt the empirical justification for the use of a sparse random projection and ridge regression approach was sufficient. In future work, we intend to expand our testing regimen to see if the empirical advantage holds in a wider range of cases.

### A.4 How does reliability thresholding impact our benchmark scores?

In the main analysis, we subselected from the greater pool of available neurons only those neurons with split-half reliabilities of 0.8 and above. In Figure 7, we show the impact of different degrees of thresholding on the scores for a majority of our models.

### A.5 How do we control for the influence of receptive field and visual acuity (resolution)?

The answer, in brief, is that we don’t – at least not explicitly. We rely instead on the reweighting procedure inherent to the neural regression method or on averaging across population responses in the representational similarity analysis to account for these properties implicitly. This may or may not be to our detriment. Recent work in both primates and rodents [9, 105–107] suggests that controlling for these factors explicitly (for example, by reshaping, translating or downsampling the resolution of the input) can in certain cases augment the strength and interpretability of the correspondence between the biological and artificial neural substrate. When designing the pipeline for the current study, however, we found that reducing the resolution of the input image by a factor of 2 and a factor of 4 (downsampling from [224,224] to [112,112] and [64,64]) in a representative sample of 12 trained and randomly-initialized models tended only to slightly shift the *absolute* depth of the features that best predict the neural activity, but changed neither the relative scores across models, nor the relative depths of the layers corresponding to different cortical areas. In future work, we hope to revisit this manipulation with a more representative sample of architectures and more vigilant consideration of how these manipulations interface with the input transforms required when extracting the features of pretrained models. Best practice for this interfacing remains unclear in the absence of more comprehensive empirical controls. The use of *custom models* (as in Nayebi et al. [88] and Shi et al. [97]), on the other hand, does not carry with it the same concerns as the use of pretrained models, and seems a promising path forward for probing the effects of input manipulations directly.

### A.6 Does training or architecture matter more for better prediction?

The range in scores between the best and worst performing model architecture trained on ImageNet is 0.209 to 0.121 (0.088) for the SRP-Ridge Max and 0.458 to −0.117 (0.575) for the RSA Max metric (excluding the two normalization-free architectures that produced negative scores and are otherwise significant outliers, the range is more like 0.458 to 0.347 (0.111)); the range between the best and worst performing model in Taskonomy is 0.190 to 0.126 (0.064) for the SRP-Ridge Max and 0.440 to 0.331 (0.109) for RSA Max. These results suggest that neither architecture nor task has a statically meaningful edge in augmenting neural predictivity. Moreover, it’s worth noting that the rankings in terms of both task and architecture show only minor differences between the best-performing and the next-best-performing models – with no major increases in performance evident across different design design and training choices, only marginal relative gains.

### A.7 Addendum: What kinds of architectures best predict rodent visual cortex?

Figure 8 provides the rankings for all ‘model zoology’ architectures in our survey.

### A.8 Addendum: What kinds of architectures best predict rodent visual cortex?

Figure 9 provides the rankings for all taskonomy encoders, clustered by task transfer affinity.

### A.9 Addendum: How do self-supervised models compare to category-supervised models?

Figure 10 provides the rankings for all the self-supervised models in our survey, along with reference points for architecture-matched category-supervised models.

### A.10 Addendum: How well do non-neural network baselines predict rodent visual cortex?

The averages we report in the main analysis with respect to two of our non-neural network baselines (gabors and GISTPCs) do obscure differences across cortical area (and cortical layer). One of the most conspicuous examples is in the case of the scores for our bank of gabors in layer 6 of primary visual cortex (VISp). At *r* = 0.165, (according to the SRP-Ridge) metric, the predictions produced by these features are competitive with the those of both trained and random deep net models.

In future work, we intend to expand our set of non-neural network baselines, incorporating perhaps the larger, more robust set of handcrafted Gabors used in custom CNN models like VOneNet [108].

### A.11 Can we visualize functional specialization even if we can’t characterize it?

In the main analysis, we show that (in general) taskonomy models fail to differentiate one cortical area from another in terms of predictivity. Here, we demonstrate this is not necessarily because these areas aren’t functionally specialized. Comparing neurons *within* a cortical site to neurons *between* using both a pure correlation-based measure, and the same style of neural regression we use in the main analysis, we show that neurons from the same site are categorically better predictors than neurons from other sites, connoting a stronger representational correspondence to neurons in anatomical proximity. Results from this analysis are available in Figure 11.

### A.12 Are there differences in model predictions across *genetic cre line*?

In the main analysis, we aggregate neurons by anatomical region (cortical area); another method of aggregating neurons is by genetic cre line. Aggregating in this way changes the overall focus of the benchmarking, from asking ‘where’ certain models fare best in predicting visual cortical activity to ‘with what cell types’. The kinds of representational idiosyncrasies that characterize different cell types are beyond the scope of this paper. Nevertheless, as a sampler for those interested, aggregate Taskonomy scores across cre line are provided in Figure 12.

### A.13 Glossary of Visual Cortical Areas in Mouse Brain

Reproduced in Figure 13 is a glossary of visual cortical areas in the mouse brain. More information about the Allen Brain Observatory visual coding dataset may be found at their website: http://observatory.brain-map.org/visualcoding

### A.14 Taskonomy Task Definitions

Reproduced in Figure 14 are Taskonomy’s official definitions of its constituent tasks. Further information is available at their website: http://taskonomy.stanford.edu

**Figure 14:**
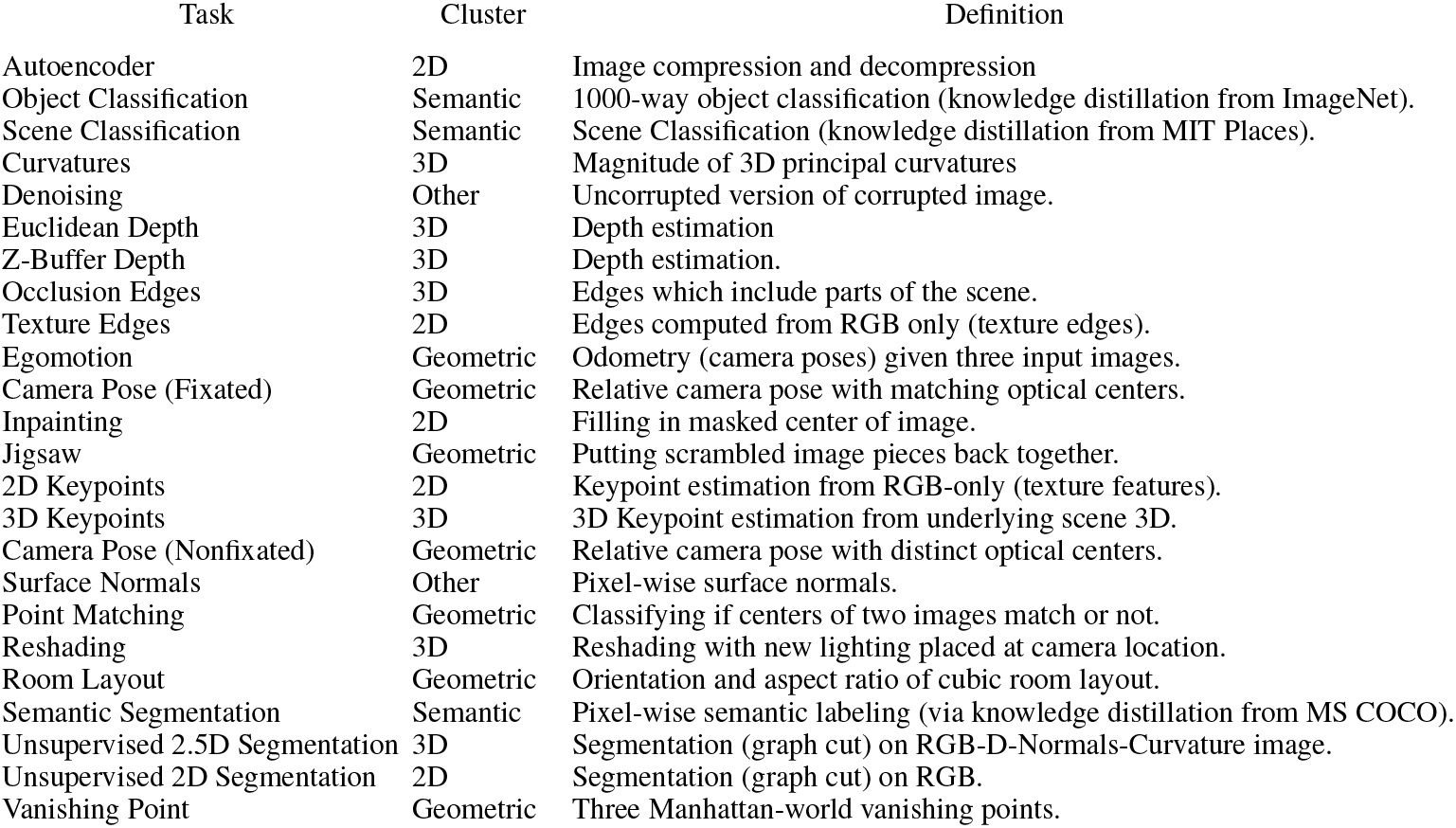
Task definitions and affinity clusters provided by Taskonomy [53].

## Footnotes

1 Available with a non-commercial license under the Allen Institute terms of use: http://www.alleninstitute.org/legal/terms-use/

2 More details are available in the whitepapers released with the observatory data: http://observatory.brain-map.org/visualcoding/transgenic

3 Note that in cases where the dimensionality of features is less than the number of projections suggested by the lemma, sparse random projections will actually upsample the feature space, rather than downsample it.

4 The use of generalized cross-validation is particularly beneficial in datasets with fewer probe images, where *k*-fold cross-validation means losing a significant degree of information in each fit.

## References

[1] Daniel LK Yamins, Ha Hong, Charles F Cadieu, Ethan A Solomon, Darren Seibert, and James J DiCarlo. Performance-optimized hierarchical models predict neural responses in higher visual cortex. Proceedings of the National Academy of Sciences, 111(23):8619–8624, 2014.

[2] Daniel LK Yamins and James J DiCarlo. Using goal-driven deep learning models to understand sensory cortex. Nature neuroscience, 19(3):356, 2016.

[3] Santiago A Cadena, George H Denfield, Edgar Y Walker, Leon A Gatys, Andreas S Tolias, Matthias Bethge, and Alexander S Ecker. Deep convolutional models improve predictions of macaque v1 responses to natural images. PLoS computational biology, 15(4):e1006897, 2019.

[4] Martin Schrimpf, Jonas Kubilius, Ha Hong, Najib J. Majaj, Rishi Rajalingham, Elias B. Issa, Kohitij Kar, Pouya Bashivan, Jonathan Prescott-Roy, Franziska Geiger, Kailyn Schmidt, Daniel L. K. Yamins, and James J. DiCarlo. Brain-score: Which artificial neural network for object recognition is most brain-like? bioRxiv preprint, 2018.

[5] Rishi Rajalingham, E.B. Issa, P. Bashivan, K. Kar, K. Schmidt, and J.J. DiCarlo. Large-scale, high-resolution comparison of the core visual object recognition behavior of humans, monkeys, and state-of-the-art deep artificial neural networks. The Journal of Neuroscience, 38(33): 7255–7269. doi: 10.1523/JNEUROSCI.0388-18.2018. URL https://doi.org/10.1523/JNEUROSCI.0388-18.2018.

[6] Pouya Bashivan, Kohitij Kar, and James J DiCarlo. Neural population control via deep image synthesis. Science, 364(6439), 2019.

[7] Jianghong Shi, Eric Shea-Brown, and Michael Buice. Comparison against task driven artificial neural networks reveals functional properties in mouse visual cortex. In Advances in Neural Information Processing Systems, pages 5765–5775, 2019.

[8] Kasper Vinken and Hans Op de Beeck. Deep neural networks point to mid-level complexity of rodent object vision. bioRxiv, 2020.

[9] S. A. Cadena, F. H. Sinz, T. Muhammad, E. Froudarakis, E. Cobos, E. Y. Walker, J. Reimer, M. Bethge, A. Tolias, and A. S. Ecker. How well do deep neural networks trained on object recognition characterize the mouse visual system? NeurIPS Neuro AI Workshop, 2019.

[10] Leslie M Kay. Olfactory coding: random scents make sense. Current Biology, 21(22): R928–R929, 2011.

[11] Davide Zoccolan, Nadja Oertelt, James J DiCarlo, and David D Cox. A rodent model for the study of invariant visual object recognition. Proceedings of the National Academy of Sciences, 106(21):8748–8753, 2009.

[12] Matthew B Broschard, Jangjin Kim, Bradley C Love, and John H Freeman. Category learning in rodents using touchscreen-based tasks. Genes, Brain and Behavior, 20(1):e12665, 2021.

[13] Jennifer L Hoy, Iryna Yavorska, Michael Wehr, and Cristopher M Niell. Vision drives accurate approach behavior during prey capture in laboratory mice. Current Biology, 26(22):3046–3052, 2016.

[14] Saskia EJ de Vries, Jerome A Lecoq, Michael A Buice, Peter A Groblewski, Gabriel K Ocker, Michael Oliver, David Feng, Nicholas Cain, Peter Ledochowitsch, Daniel Millman, et al. A large-scale standardized physiological survey reveals functional organization of the mouse visual cortex. Nature Neuroscience, 23(1):138–151, 2020.

[15] Simon Musall, Matthew T Kaufman, Ashley L Juavinett, Steven Gluf, and Anne K Churchland. Single-trial neural dynamics are dominated by richly varied movements. Nature neuroscience, 22(10):1677–1686, 2019.

[16] Adam Paszke, Sam Gross, Francisco Massa, Adam Lerer, James Bradbury, Gregory Chanan, Trevor Killeen, Zeming Lin, Natalia Gimelshein, Luca Antiga, Alban Desmaison, Andreas Kopf, Edward Yang, Zachary DeVito, Martin Raison, Alykhan Tejani, Sasank Chilamkurthy, Benoit Steiner, Lu Fang, Junjie Bai, and Soumith Chintala. Pytorch: An imperative style, high-performance deep learning library. In Advances in Neural Information Processing Systems 32, pages 8024–8035. 2019.

[17] Ross Wightman. Pytorch image models. https://github.com/rwightman/pytorch-image-models, 2019.

[18] Alex Krizhevsky, Ilya Sutskever, and Geoffrey E Hinton. ImageNet classification with deep convolutional neural networks. In Advances in Neural Information Processing Systems, pages 1097–1105, 2012.

[19] Gao Huang, Zhuang Liu, Laurens Van Der Maaten, and Kilian Q Weinberger. Densely connected convolutional networks. In Conference on Computer Vision and Pattern Recognition, pages 4700–4708, 2017.

[20] Christian Szegedy, Wei Liu, Yangqing Jia, Pierre Sermanet, Scott Reed, Dragomir Anguelov, Dumitru Erhan, Vincent Vanhoucke, and Andrew Rabinovich. Going deeper with convolutions. In Conference on Computer Vision and Pattern Recognition, pages 1–9, 2015.

[21] Mingxing Tan, Bo Chen, Ruoming Pang, Vijay Vasudevan, Mark Sandler, Andrew Howard, and Quoc V Le. Mnasnet: Platform-aware neural architecture search for mobile. In Conference on Computer Vision and Pattern Recognition, pages 2820–2828, 2019.

[22] Mark Sandler, Andrew Howard, Menglong Zhu, Andrey Zhmoginov, and Liang-Chieh Chen. Mobilenetv2: Inverted residuals and linear bottlenecks. In Conference on Computer Vision and Pattern Recognition, pages 4510–4520, 2018.

[23] Kaiming He, Xiangyu Zhang, Shaoqing Ren, and Jian Sun. Deep residual learning for image recognition. In Conference on Computer Vision and Pattern Recognition, pages 770–778, 2016.

[24] Saining Xie, Ross Girshick, Piotr Dollár, Zhuowen Tu, and Kaiming He. Aggregated residual transformations for deep neural networks. In Conference on Computer Vision and Pattern Recognition, pages 1492–1500, 2017.

[25] Xiangyu Zhang, Xinyu Zhou, Mengxiao Lin, and Jian Sun. Shufflenet: An extremely efficient convolutional neural network for mobile devices. In Conference on Computer Vision and Pattern Recognition, pages 6848–6856, 2018.

[26] Karen Simonyan and Andrew Zisserman. Very deep convolutional networks for large-scale image recognition. 2014. https://arxiv.org/abs/1409.1556.

[27] Sergey Zagoruyko and Nikos Komodakis. Wide residual networks. 2016. https://arxiv.org/abs/1605.07146.

[28] Forrest N Iandola, Song Han, Matthew W Moskewicz, Khalid Ashraf, William J Dally, and Kurt Keutzer. Squeezenet: Alexnet-level accuracy with 50x fewer parameters and¡ 0.5 mb model size. 2016. https://arxiv.org/abs/1602.07360.

[29] Alexey Bochkovskiy, Chien-Yao Wang, and Hong-Yuan Mark Liao. Yolov4: Optimal speed and accuracy of object detection. CVPR, 2020. https://arxiv.org/abs/2004.10934.

[30] Christian Szegedy, Sergey Ioffe, Vincent Vanhoucke, and Alex Alemi. Inception-v4, inception-resnet and the impact of residual connections on learning. AAAI, 2017. https://arxiv.org/abs/1602.07261.

[31] Mingxing Tan and Quoc V. Le. Efficientnet: Rethinking model scaling for convolutional neural networks. ICML, 2019. https://arxiv.org/abs/1905.11946.

[32] Fisher Yu, Dequan Wang, Evan Shelhamer, and Trevor Darrell. Deep layer aggregation. CVPR, 2018. https://arxiv.org/abs/1707.06484.

[33] Qilong Wang, Banggu Wu, Pengfei Zhu, Peihua Li, Wangmeng Zuo, and Qinghua Hu. Eca-net: Efficient channel attention for deep convolutional neural networks. CVPR, 2020. https://arxiv.org/abs/1910.03151.

[34] Bichen Wu, Xiaoliang Dai, Peizhao Zhang, Yanghan Wang, Fei Sun, Yiming Wu, Yuandong Tian, Peter Vajda, Yangqing Jia, and Kurt Keutzer. Fbnet: Hardware-aware efficient convnet design via differentiable neural architecture search, 2019. https://arxiv.org/abs/1812.03443.

[35] Mingxing Tan and Quoc V. Le. Mixconv: Mixed depthwise convolutional kernels. BMVC, 2019. https://arxiv.org/abs/1907.09595.

[36] Alexey Dosovitskiy, Lucas Beyer, Alexander Kolesnikov, Dirk Weissenborn, Xiaohua Zhai, Thomas Unterthiner, Mostafa Dehghani, Matthias Minderer, Georg Heigold, Sylvain Gelly, Jakob Uszkoreit, and Neil Houlsby. An image is worth 16×16 words: Transformers for image recognition at scale. CVPR, 2020. https://arxiv.org/abs/2010.11929.

[37] Xiang Li, Wenhai Wang, Xiaolin Hu, and Jian Yang. Selective kernel networks. CVPR, 2019. https://arxiv.org/abs/1903.06586.

[38] Chenxi Liu, Barret Zoph, Maxim Neumann, Jonathon Shlens, Wei Hua, Li-Jia Li, Li Fei-Fei, Alan Yuille, Jonathan Huang, and Kevin Murphy. Progressive neural architecture search. ECCV, 2018. https://arxiv.org/abs/1712.00559.

[39] Jie Hu, Li Shen, Samuel Albanie, Gang Sun, and Enhua Wu. Squeeze-and-excitation networks. CVPR, 2019. https://arxiv.org/abs/1709.01507.

[40] Ze Liu, Yutong Lin, Yue Cao, Han Hu, Yixuan Wei, Zheng Zhang, Stephen Lin, and Baining Guo. Swin transformer: Hierarchical vision transformer using shifted windows, 2021. https://arxiv.org/abs/2103.14030.

[41] François Chollet. Xception: Deep learning with depthwise separable convolutions, 2017. https://arxiv.org/abs/1610.02357.

[42] Niv Nayman, Yonathan Aflalo, Asaf Noy, and Lihi Zelnik-Manor. Hardcore-nas: Hard constrained differentiable neural architecture search, 2021. https://arxiv.org/abs/2102.11646.

[43] Stéphane d’Ascoli, Hugo Touvron, Matthew Leavitt, Ari Morcos, Giulio Biroli, and Levent Sagun. Convit: Improving vision transformers with soft convolutional inductive biases, 2021. https://arxiv.org/abs/2103.10697.

[44] Weijian Xu, Yifan Xu, Tyler Chang, and Zhuowen Tu. Co-scale conv-attentional image transformers. ICCV, 2021. https://arxiv.org/abs/2104.06399.

[45] Kai Han, Yunhe Wang, Qi Tian, Jianyuan Guo, Chunjing Xu, and Chang Xu. Ghostnet: More features from cheap operations. CVPR, 2020. https://arxiv.org/abs/1911.11907.

[46] Ben Graham, Alaaeldin El-Nouby, Hugo Touvron, Pierre Stock, Armand Joulin, Hervé Jégou, and Matthijs Douze. Levit: a vision transformer in convnet’s clothing for faster inference, 2021. https://arxiv.org/abs/2104.01136.

[47] Ilya Tolstikhin, Neil Houlsby, Alexander Kolesnikov, Lucas Beyer, Xiaohua Zhai, Thomas Unterthiner, Jessica Yung, Andreas Steiner, Daniel Keysers, Jakob Uszkoreit, Mario Lucic, and Alexey Dosovitskiy. Mlp-mixer: An all-mlp architecture for vision, 2021. https://arxiv.org/abs/2105.01601.

[48] Andrew Howard, Mark Sandler, Grace Chu, Liang-Chieh Chen, Bo Chen, Mingxing Tan, Weijun Wang, Yukun Zhu, Ruoming Pang, Vijay Vasudevan, Quoc V. Le, and Hartwig Adam. Searching for mobilenetv3. ICCV, 2019. https://arxiv.org/abs/1905.02244.

[49] Andrew Brock, Soham De, Samuel L. Smith, and Karen Simonyan. High-performance large-scale image recognition without normalization, 2021. https://arxiv.org/abs/2102.06171.

[50] Andrew Brock, Soham De, and Samuel L. Smith. Characterizing signal propagation to close the performance gap in unnormalized resnets. ICLR, 2021. https://arxiv.org/abs/2101.08692.

[51] Xiaohan Ding, Xiangyu Zhang, Ningning Ma, Jungong Han, Guiguang Ding, and Jian Sun. Repvgg: Making vgg-style convnets great again. CVPR, 2021. https://arxiv.org/abs/2101.03697.

[52] Hugo Touvron, Piotr Bojanowski, Mathilde Caron, Matthieu Cord, Alaaeldin El-Nouby, Edouard Grave, Armand Joulin, Gabriel Synnaeve, Jakob Verbeek, and Hervé Jégou. Resmlp: Feedforward networks for image classification with data-efficient training, 2021. https://arxiv.org/abs/2105.03404.

[53] Amir R Zamir, Alexander Sax, William Shen, Leonidas J Guibas, Jitendra Malik, and Silvio Savarese. Taskonomy: Disentangling task transfer learning. In Proceedings of the IEEE Conference on Computer Vision and Pattern Recognition, pages 3712–3722, 2018.

[54] Alexander Sax, Jeffrey O Zhang, Bradley Emi, Amir Zamir, Silvio Savarese, Leonidas Guibas, and Jitendra Malik. Learning to navigate using mid-level visual priors. CoRL, 2019. https://arxiv.org/abs/1912.11121.

[55] Aria Wang, Michael Tarr, and Leila Wehbe. Neural taskonomy: Inferring the similarity of task-derived representations from brain activity. In Advances in Neural Information Processing Systems, pages 15475–15485, 2019.

[56] Zhirong Wu, Yuanjun Xiong, Stella X Yu, and Dahua Lin. Unsupervised feature learning via non-parametric instance discrimination. In Proceedings of the IEEE conference on computer vision and pattern recognition, pages 3733–3742, 2018.

[57] Ting Chen, Simon Kornblith, Mohammad Norouzi, and Geoffrey Hinton. A simple framework for contrastive learning of visual representations. In International conference on machine learning, pages 1597–1607. PMLR, 2020.

[58] Chengxu Zhuang, Siming Yan, Aran Nayebi, Martin Schrimpf, Michael C Frank, James J DiCarlo, and Daniel LK Yamins. Unsupervised neural network models of the ventral visual stream. Proceedings of the National Academy of Sciences, 118(3), 2021.

[59] Talia Konkle and George A Alvarez. Beyond category-supervision: instance-level contrastive learning models predict human visual system responses to objects. bioRxiv, 2021.

[60] Priya Goyal, Quentin Duval, Jeremy Reizenstein, Matthew Leavitt, Min Xu, Benjamin Lefaudeux, Mannat Singh, Vinicius Reis, Mathilde Caron, Piotr Bojanowski, Armand Joulin, and Ishan Misra. Vissl. https://github.com/facebookresearch/vissl, 2021.

[61] Mathilde Caron, Piotr Bojanowski, Armand Joulin, and Matthijs Douze. Deep clustering for unsupervised learning of visual features. In Proceedings of the European Conference on Computer Vision (ECCV), pages 132–149, 2018.

[62] Kaiming He, Haoqi Fan, Yuxin Wu, Saining Xie, and Ross Girshick. Momentum contrast for unsupervised visual representation learning. In Proceedings of the IEEE/CVF Conference on Computer Vision and Pattern Recognition, pages 9729–9738, 2020.

[63] Mathilde Caron, Ishan Misra, Julien Mairal, Priya Goyal, Piotr Bojanowski, and Armand Joulin. Unsupervised learning of visual features by contrasting cluster assignments. NeurIPS, 2020. https://arxiv.org/abs/2006.09882.

[64] Mathilde Caron, Hugo Touvron, Ishan Misra, Hervé Jégou, Julien Mairal, Piotr Bojanowski, and Armand Joulin. Emerging properties in self-supervised vision transformers. In Proceedings of the International Conference on Computer Vision (ICCV), 2021.

[65] Jure Zbontar, Li Jing, Ishan Misra, Yann LeCun, and Stéphane Deny. Barlow twins: Selfsupervised learning via redundancy reduction. ICML, 2021. https://arxiv.org/abs/2103.03230.

[66] Nikolaus Kriegeskorte, Marieke Mur, Douglas A Ruff, Roozbeh Kiani, Jerzy Bodurka, Hossein Esteky, Keiji Tanaka, and Peter A Bandettini. Matching categorical object representations in inferior temporal cortex of man and monkey. Neuron, 60(6):1126–1141, 2008.

[67] Umut Güçlü and Marcel AJ van Gerven. Increasingly complex representations of natural movies across the dorsal stream are shared between subjects. NeuroImage, 145:329–336, 2017.

[68] Matteo Carandini, Jonathan B Demb, Valerio Mante, David J Tolhurst, Yang Dan, Bruno A Olshausen, Jack L Gallant, and Nicole C Rust. Do we know what the early visual system does? Journal of Neuroscience, 25(46):10577–10597, 2005.

[69] N. Kriegeskorte, M. Mur, D.A. Ruff, R. Kiani, J. Bodurka, H. Esteky, and P.A. Bandettini. Matching categorical object representations in inferior temporal cortex of man and monkey. Neuron, 60(6):1126–1141. doi: 10.1016/j.neuron.2008.10.043. URL https://doi.org/10.1016/j.neuron.2008.10.043.

[70] F. Pedregosa, G. Varoquaux, A. Gramfort, V. Michel, B. Thirion, O. Grisel, M. Blondel, P. Prettenhofer, R. Weiss, V. Dubourg, J. Vanderplas, A. Passos, D. Cournapeau, M. Brucher, M. Perrot, and E. Duchesnay. Scikit-learn: Machine learning in Python. Journal of Machine Learning Research, 12:2825–2830, 2011.

[71] Ali Rahimi, Benjamin Recht, et al. Random features for large-scale kernel machines. In NIPS, volume 3, page 5. Citeseer, 2007.

[72] Aude Oliva and Antonio Torralba. Modeling the shape of the scene: A holistic representation of the spatial envelope. International journal of computer vision, 42(3):145–175, 2001.

[73] Kamila Maria Jozwik, Martin Schrimpf, Nancy Kanwisher, and James J DiCarlo. To find better neural network models of human vision, find better neural network models of primate vision. BioRxiv, page 688390, 2019.

[74] D.J. Felleman and D.C. Van Essen. Distributed hierarchical processing in the primate cerebral cortex. Cerebral Cortex, 1(1):1–47.

[75] J.J. DiCarlo, D. Zoccolan, and N.C. Rust. How does the brain solve visual object recognition? Neuron, 73(3):415–434. doi: 10.1016/j.neuron.2012.01.010. URL https://doi.org/10.1016/j.neuron.2012.01.010.

[76] Nicole C Rust and James J DiCarlo. Selectivity and tolerance (“invariance”) both increase as visual information propagates from cortical area v4 to it. Journal of Neuroscience, 30(39): 12978–12995, 2010.

[77] Quanxin Wang, Enquan Gao, and Andreas Burkhalter. Gateways of ventral and dorsal streams in mouse visual cortex. Journal of Neuroscience, 31(5):1905–1918, 2011.

[78] Quanxin Wang, Olaf Sporns, and Andreas Burkhalter. Network analysis of corticocortical connections reveals ventral and dorsal processing streams in mouse visual cortex. Journal of Neuroscience, 32(13):4386–4399, 2012.

[79] Rinaldo D D’Souza, Quanxin Wang, Weiqing Ji, Andrew M Meier, Henry Kennedy, Kenneth Knoblauch, and Andreas Burkhalter. Canonical and noncanonical features of the mouse visual cortical hierarchy. bioRxiv, 2020.

[80] Joshua H Siegle, Xiaoxuan Jia, Séverine Durand, Sam Gale, Corbett Bennett, Nile Graddis, Greggory Heller, Tamina K Ramirez, Hannah Choi, Jennifer A Luviano, et al. Survey of spiking in the mouse visual system reveals functional hierarchy. Nature, 592(7852):86–92, 2021.

[81] Mark L Andermann, Aaron M Kerlin, Demetris K Roumis, Lindsey L Glickfeld, and R Clay Reid. Functional specialization of mouse higher visual cortical areas. Neuron, 72(6):1025–1039, 2011.

[82] Joshua H Siegle, Peter Ledochowitsch, Xiaoxuan Jia, Daniel J Millman, Gabriel K Ocker, Shiella Caldejon, Linzy Casal, Andy Cho, Daniel J Denman, Séverine Durand, et al. Reconciling functional differences in populations of neurons recorded with two-photon imaging and electrophysiology. Elife, 10:e69068, 2021.

[83] Julie A Harris, Stefan Mihalas, Karla E Hirokawa, Jennifer D Whitesell, Hannah Choi, Amy Bernard, Phillip Bohn, Shiella Caldejon, Linzy Casal, Andrew Cho, et al. Hierarchical organization of cortical and thalamic connectivity. Nature, 575(7781):195–202, 2019.

[84] Nancy Kanwisher. Functional specificity in the human brain: a window into the functional architecture of the mind. Proceedings of the National Academy of Sciences, 107(25):11163–11170, 2010.

[85] Kalanit Grill-Spector and Kevin S Weiner. The functional architecture of the ventral temporal cortex and its role in categorization. Nature Reviews Neuroscience, 15(8):536–548, 2014.

[86] Dwight J Kravitz, Kadharbatcha S Saleem, Chris I Baker, and Mortimer Mishkin. A new neural framework for visuospatial processing. Nature Reviews Neuroscience, 12(4):217–230, 2011.

[87] Jörn Diedrichsen and Nikolaus Kriegeskorte. Representational models: A common framework for understanding encoding, pattern-component, and representational-similarity analysis. PLoS computational biology, 13(4):e1005508, 2017.

[88] Aran Nayebi, Nathan CL Kong, Chengxu Zhuang, Justin L Gardner, Anthony M Norcia, and Daniel LK Yamins. Unsupervised models of mouse visual cortex. bioRxiv, 2021.

[89] K. Kar, J. Kubilius, K. Schmidt, E.B. Issa, and J.J. DiCarlo. Evidence that recurrent circuits are critical to the ventral stream’s execution of core object recognition behavior. Nature Neuroscience. doi: 10.1038/s41593-019-0392-5. URL https://doi.org/10.1038/s41593-019-0392-5.

[90] Jonas Kubilius, Martin Schrimpf, Ha Hong, Najib J. Majaj, Rishi Rajalingham, Elias B. Issa, Kohitij Kar, Pouya Bashivan, Jonathan Prescott-Roy, Kailyn Schmidt, Aran Nayebi, Daniel Bear, Daniel L. K. Yamins, and James J. DiCarlo. Brain-Like Object Recognition with High-Performing Shallow Recurrent ANNs. In Neural Information Processing Systems (NeurIPS), pages 12785–-12796. Curran Associates, Inc., 2019.

[91] Ivan Ustyuzhaninov, Santiago A Cadena, Emmanouil Froudarakis, Paul G Fahey, Edgar Y Walker, Erick Cobos, Jacob Reimer, Fabian H Sinz, Andreas S Tolias, Matthias Bethge, et al. Rotation-invariant clustering of neuronal responses in primary visual cortex. In International Conference on Learning Representations, 2019.

[92] Sen Wang, Ronald Clark, Hongkai Wen, and Niki Trigoni. Deepvo: Towards end-to-end visual odometry with deep recurrent convolutional neural networks. 2017 IEEE International Conference on Robotics and Automation (ICRA), May 2017. doi: 10.1109/icra.2017.7989236. URL http://dx.doi.org/10.1109/ICRA.2017.7989236.

[93] GT Prusky and RM Douglas. Characterization of mouse cortical spatial vision. Vision research, 44(28):3411–3418, 2004.

[94] Chengxu Zhuang, Jonas Kubilius, Mitra JZ Hartmann, and Daniel L Yamins. Toward goal-driven neural network models for the rodent whisker-trigeminal system. In Advances in Neural Information Processing Systems, pages 2555–2565, 2017.

[95] Anthony M Zador. A critique of pure learning and what artificial neural networks can learn from animal brains. Nature communications, 10(1):1–7, 2019.

[96] Josh Merel, Diego Aldarondo, Jesse Marshall, Yuval Tassa, Greg Wayne, and Bence Ölveczky. Deep neuroethology of a virtual rodent. 2019. https://arxiv.org/abs/1911.09451.

[97] Jianghong Shi, Bryan Tripp, Eric Shea-Brown, Stefan Mihalas, and Michael Buice. Cnn mousenet: A biologically constrained convolutional neural network model for mouse visual cortex. bioRxiv, 2021.

[98] Seyed-Mahdi Khaligh-Razavi and Nikolaus Kriegeskorte. Deep supervised, but not unsupervised, models may explain it cortical representation. PLoS computational biology, 10(11), 2014.

[99] David A Klindt, Alexander S Ecker, Thomas Euler, and Matthias Bethge. Neural system identification for large populations separating” what” and” where”. 2017. https://arxiv.org/abs/1711.02653.

[100] Ari Morcos, Maithra Raghu, and Samy Bengio. Insights on representational similarity in neural networks with canonical correlation. In Advances in Neural Information Processing Systems, pages 5727–5736, 2018.

[101] Simon Kornblith, Mohammad Norouzi, Honglak Lee, and Geoffrey Hinton. Similarity of neural network representations revisited. ICML, 2019. https://arxiv.org/abs/1905.00414.

[102] Natalia Y Bilenko and Jack L Gallant. Pyrcca: regularized kernel canonical correlation analysis in python and its applications to neuroimaging. Frontiers in neuroinformatics, 10:49, 2016.

[103] Ryan M Rifkin and Ross A Lippert. Notes on regularized least squares. 2007.

[104] Leyla Tarhan and Talia Konkle. Reliability-based voxel selection. NeuroImage, 207:116350, 2020.

[105] S.A. Cadena, G.H. Denfield, E.Y. Walker, L.A. Gatys, A.S. Tolias, M. Bethge, and A.S. Ecker. Deep convolutional models improve predictions of macaque v1 responses to natural images author summary. PLoS Computational Biology, 1–28. doi: 10.12751/g-node.2e31e3. URL https://doi.org/10.12751/g-node.2e31e3.

[106] Edgar Y Walker, Fabian H Sinz, Erick Cobos, Taliah Muhammad, Emmanouil Froudarakis, Paul G Fahey, Alexander S Ecker, Jacob Reimer, Xaq Pitkow, and Andreas S Tolias. Inception loops discover what excites neurons most using deep predictive models. Nature neuroscience, 22(12):2060–2065, 2019.

[107] Tiago Marques, Martin Schrimpf, and James J DiCarlo. Multi-scale hierarchical neural network models that bridge from single neurons in the primate primary visual cortex to object recognition behavior. bioRxiv, 2021.

[108] Joel Dapello, Tiago Marques, Martin Schrimpf, Franziska Geiger, David D Cox, and James J DiCarlo. Simulating a primary visual cortex at the front of cnns improves robustness to image perturbations. BioRxiv, 2020.

